# Structural features within precursor microRNA-20a regulate Dicer-TRBP processing

**DOI:** 10.1101/2025.05.07.652689

**Authors:** Yaping Liu, Cade T. Harkner, Megan N. Westwood, Aldrex Munsayac, Sarah C. Keane

## Abstract

MicroRNAs (miRNAs) are small non-coding RNAs that post-transcriptionally regulate gene expression of target messenger (m) RNAs. To maintain proper miRNA expression levels, the enzymatic processing of primary and precursor miRNA elements must be strictly controlled. However, the molecular determinants underlying this strict regulation of miRNA biogenesis are not fully understood. Here, we determined the solution structure of pre-miR-20a, an oncogenic miRNA and component of the oncomiR-1 cluster, using nuclear magnetic spectroscopy (NMR) spectroscopy and small angle X-ray scattering (SAXS). Our structural studies informed on key secondary structure elements of pre-miR-20a which may control its enzymatic processing, namely a flexible apical loop and single-nucleotide bulge near the dicing site. We found that alternative conformations within pre-miR-20a’s apical loop function to self-regulate its Dicer-TRBP processing, and that a single nucleotide bulge at the -5 position from the 5′-cleavage site is critical for efficient processing. We additionally found that a disease-related single-nucleotide polymorphism in pre-miR-20a, predicted to disrupt the structure near the dicing site, resulted in reduced processing. These results further our structural understanding of the oncomiR-1 cluster and show how transient RNA conformers can function to self-regulate maturation.

## Introduction

MicroRNAs (miRNAs) are highly-conserved, short (∼22 nucleotide), single-stranded non-coding RNAs that, in coordination with the Argonaut (Ago) protein, function to regulate gene expression through interactions with the 3ʹ-untranslated regions (UTRs) of messenger RNAs (mRNAs).^1–4^ Mature miRNAs are the products of sequential enzymatic processing events. In the nucleus, most primary miRNAs (pri-miRNAs) are transcribed by RNA polymerase II and enzymatically cleaved into a precursor microRNA (pre-miRNA) by the microprocessor complex (MP), primarily composed of the RNase III enzyme Drosha and partner protein Di-George Critical Region 8 (DGCR8), along with various accessory proteins.^5–10^ The pre-miRNA hairpin is then exported into the cytoplasm by Exportin 5 (XPO5) and Ran-GTPase.^11–13^ In the cytoplasm, another RNase III enzyme Dicer in complex with the transactivation response element RNA-binding protein (TRBP) cleaves the apical loop from the pre-miRNA, generating a mature miRNA duplex.^14–16^ One strand of the mature miRNA duplex is then loaded into Ago, generating the RNA-induced silencing complex (RISC), which interacts with and degrades or prevents translation of mRNA targets.^3,17–19^

To maintain proper miRNA expression levels, the enzymatic processing of pri- and pre-miRNA elements must be strictly controlled. Critically, both RNA-protein interactions^20–24^ and RNA structures and sequence motifs can function to regulate miRNA biogenesis at both MP and Dicer cleavage events.^25–32^ For example, proteins such as hnRNPA1, RBFOX, and LIN28 bind to pri- or pre-miRNAs and remodel RNA structure or recruit accessory protein factors and/or MP.^22,33,34^ Enzymatic processing is dependent on certain RNA sequence elements for efficient processing, such as UGU^35,36^ and fUN/UG motifs^35,37,38^ on pri-miRNAs for DGCR8 and Drosha recognition, respectively. Processing efficiency can also be influenced by innate structural properties of the pri- and pre-miRNAs, such as the positioning and size of internal and apical loops.^25–32^ A growing body of research has identified innate structural dynamics of pre-miRNAs as key regulatory factors in processing. Previous studies have revealed that dynamics in the apical loop junction of pre-miR-31 near the dicing site promotes distinct favorable interactions,^25^ and that pH-dependent structural rearrangements of pre-miR-21 can significantly improve Dicer processing.^30^

In this work we investigate the structure of pre-miR-20a, a component of the miR-17∼92a (oncomiR-1) pri-miRNA cluster, and the RNA features that contribute to regulation of dicing. Controlling miR-20a levels is crucial for cellular homeostasis, as dysregulation of miR-20a has been implicated in several cancers including breast, liver, lung, endometrial, colorectal, cervical, and leukemia.^39^ Here, we determined the three-dimensional structure of pre-miR-20a by NMR spectroscopy, the first structure of a pre-miRNA from the oncomiR-1 cluster. Like many pre-miRNAs, pre-miR-20a contains an imperfectly paired stem, including a pH-dependent C-A^+^ mismatch and an A-rich internal loop.^40^ Additionally, we identified structural motifs which may be important for processing efficiency, namely a single-nucleotide G bulge just below the dicing site and alternative conformations within the apical loop of pre-miR-20a. Using our NMR-derived structure to inform mutational design, we probed the effects of these structural motifs on Dicer-TRBP processing *in vitro*. We found that sparsely populated alternative conformations within the apical loop of pre-miR-20a function to negatively regulate pre-miR-20a Dicer processing, and that the position of a single nucleotide bulge on the 5ʹ-strand was critical for efficient processing. Further, we characterized the effects of U508C, a colorectal cancer-associated mutation in pre-miR-20a. This single nucleotide polymorphism (SNP) reduced Dicer processing, likely a result of significant structural perturbation near the dicing site. Our work enhances the understanding of miR-20a biogenesis and provides further evidence that in the absence of protein-binding partners, innate RNA structural dynamics can function to self-regulate pre-miRNA Dicer processing.

## Results

### Oligonucleotide controls and deuterium editing approach for obtaining chemical shift assignments of FL pre-miR-20a

To gain molecular-level insight into the regulation of pre-miR-20a processing, we initiated structural studies using solution NMR spectroscopy. Chemical shift assignments are a prerequisite for any NMR-based structural study, however, the full-length (FL) pre-miR-20a (**Fig. 1A**) is 73 nucleotides (nts) and its large size poses some technical challenges related to spectral overlap. Therefore, we employed a “divide and conquer” approach, an approach we have successfully applied to the study of other large RNAs,^25,40,41^ to facilitate initial FL pre-miR-20a resonance assignments. We divided the FL pre-miR-20a structure into four smaller, partially overlapping oligonucleotide controls (20a-frag1, -frag2, -frag3, and -frag4), each approximately 30 nts long (**Fig. S1**). Each of the four control RNAs yielded high-quality NMR data, and we successfully completed aromatic (C2, C8, C6, H2, H8, H5, H6) and ribose (H1ʹ, H2ʹ, H3ʹ) resonance assignments (**Fig. S2-S5**).

**Figure 1.**
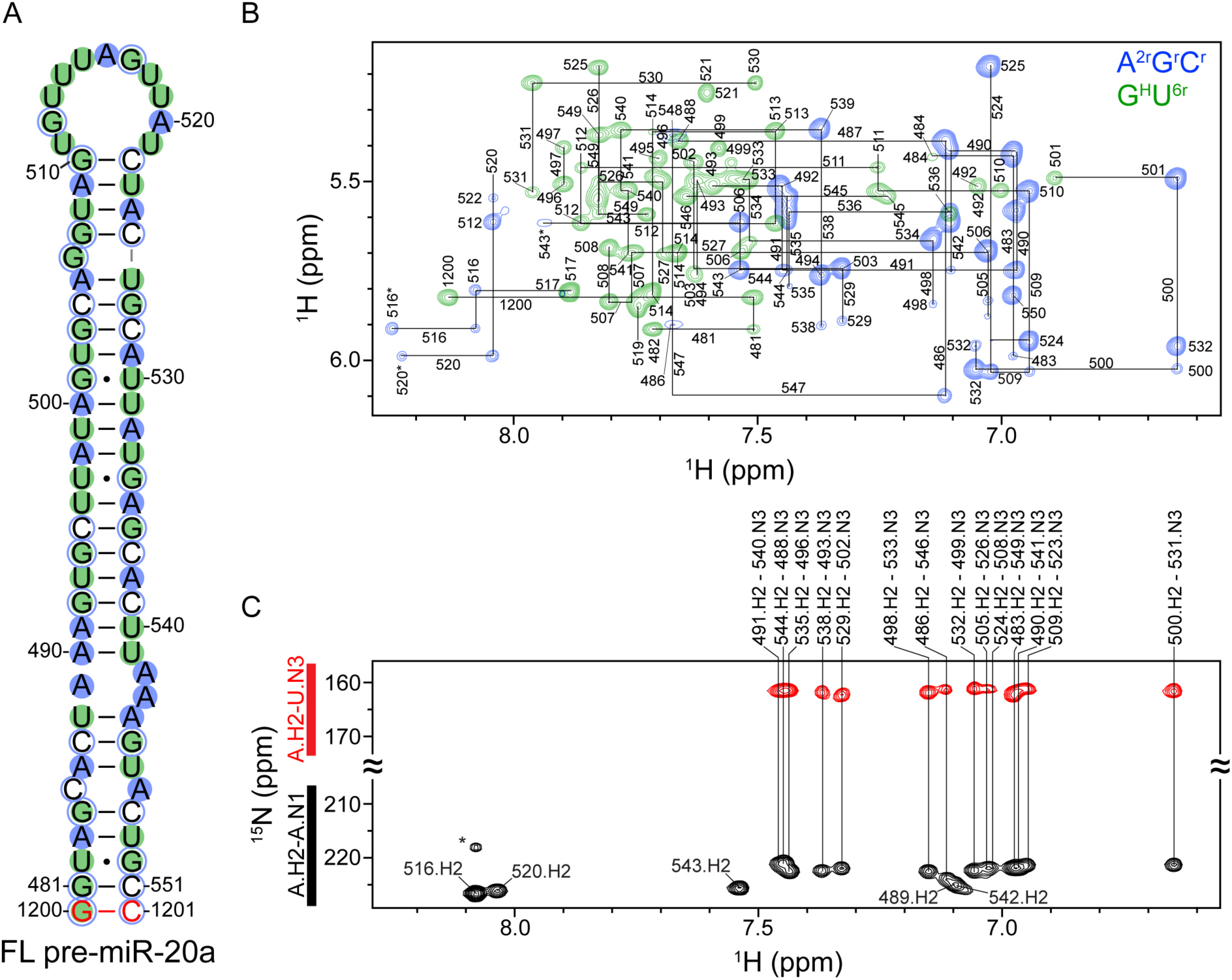
The pre-miR-20a hairpin contains an imperfectly paired stem and large apical loop. (A) Secondary structure of FL pre-miR-20a. Red nucleotides are non-native. Colored circles correspond to the deuterium labeling approach (in B). Closed circles represent nucleosides protiated on the base and ribose, open circles represent nucleosides protiated only on the ribose. (B) Overlay of A^2r^G^r^C^r^-labeled (blue) and G^H^U^6r–^labeled (green) 2D ^1^H-^1^H NOESY spectra with assignments. Assignments with an asterisk denote adenosine C8 breakthrough signals. (C) Best-selective long-range HNN-COSY spectrum identifying A-U base pairs within pre-miR-20a. black peaks are adenosine H2-N1 correlations, red peaks are adenosine H2-uracil N3 correlations. Vertical lines indicate the detection of A-U base pairs. The asterisk denotes a non-suppressed adenosine-N3 signal.

The control RNAs were designed to guide assignments of the FL pre-miR-20a RNA. However, the two-dimensional (2D) ^1^H-^1^H NOESY spectrum of pre-miR-20a exhibited significant spectral overlap and line broadening due to its large molecular size (**Fig. S6A**). These challenges made it impossible to confidently complete resonance assignments based on the chemical shifts obtained from the oligonucleotide controls. To improve the FL pre-miR-20a ^1^H-^1^H NOESY spectral quality, we used a deuterium-edited approach,^42–44^ which involved the analysis of ^1^H-^1^H NOESY spectra collected on FL pre-miR-20a RNAs prepared with different combinations of fully protiated, fully deuterated, and partially deuterated ribonucleotide triphosphates. The following FL pre-miR-20a samples were prepared: G^H^U^6r,^ A^H^C^H^U^r^, and A^2r^G^r^C^r^ (**Fig. S6B-D**). In this nomenclature, superscripts denote sites containing hydrogen atoms on a given nucleoside while all other non-exchangeable sites are deuterated (e.g. U^6r^ indicates that all uridines were protiated at the C6 and ribose carbons, A^H^ indicates the use of fully protiated adenosine). Individually, each of the deuterium-edited samples exhibited sharper line widths and less spectral crowding relative to the protiated sample (**Fig. S6**). However, the power of the deuterium-edited approach is the analysis of these spectra together (**Fig. 1B**). This combination of methods allowed for complete assignment of the nonexchangeable aromatic and ribose protons (H1ʹ, H2ʹ and H3ʹ) in FL pre-miR-20a (**Fig. S7, Table S1**).

### The secondary structure of FL pre-miR-20a contains a large apical loop and an imperfectly paired stem

To determine the secondary structure of FL pre-miR-20a, we acquired a BEST selective long-range HNN-COSY.^45^ These data support the presence of 14 A-U based pairs within the stem of FL pre-miR-20a (**Fig. 1C**). Consistent with other known and predicted pre-miRNA hairpin structures, the stem of FL pre-miR-20a contains several mismatches and internal loops and therefore does not form a perfect duplex.^40,46–49^ We observed five A.H2-A.N1 correlations that did not have corresponding A.H2-U.N3 signals, suggesting that these five adenosines are not involved in canonical A-U base pairs (**Fig. 1C**). The H2 of A547 is broadened beyond detection and represents a sixth unpaired adenosine in FL pre-miR-20a. We note that A547 may form a non-canonical base pair with C485 at low pH.^40,50,51^ Circular dichroism (CD) analysis of 20a-frag1 revealed an increased melting temperature (ΔT_m_ = 6.3 °C) at pH 5.8 relative to pH 7.5 (**Fig. S8A**). Furthermore, we observed a significant shift of the A547 C2-H2 correlation at low pH (**Fig. S8B**), indicating that A547 is protonated at the nearby N1 under low pH conditions which would promote formation of a C-A^+^ base pair. In addition to the lack of A.H2-U.N3 correlations, the downfield chemical shifts of A520.H2 and A516.H2 are consistent with unpaired adenosines, suggesting that FL pre-miR-20a contains a large apical loop.

The long-range HNN-COSY data was limited to A-U base pairs; therefore, we further interrogated the secondary structure of FL pre-miR-20a through analysis of imino proton spectra. The large size of FL pre-miR-20a precluded analysis of the imino spectrum due to spectral overlap (**Fig. S9**). We therefore used the small fragments to further define the secondary structure of regions of FL pre-miR-20a that are predicted to contain bulges, internal loops, and mismatches (**Fig. S10-12**). The imino proton spectrum collected on 20a-frag1 is consistent with the presence of nine canonical Watson-Crick-Franklin base pairs and one U•G wobble pair (**Fig. S10**). The upfield G1202.H1 chemical shift is consistent with presence of a sheared G-A base pair which forms between the first and last residues of the non-native GAGA tetraloop.^52^ Notably, the H3 of U488 is broadened beyond detection, consistent with dynamics within the 1×2 internal loop that is flanking the U488-A544 base pair.

The imino proton resonances of 20a-frag3 were identified and assigned based on the 20a-frag3 NOESY spectrum at low temperatures (**Fig. S11**). In addition to the ten signals corresponding to canonical base pairs, two peaks that correspond to the G501•U530 wobble pair, and one peak for G1216 can be assigned. We were unable to assign the imino resonances for G506 or U526 and the resonance for G507 is too broad to observe in the 2D NOESY, consistent with potential dynamics in this region. However, in the imino proton spectrum collected on 20a-frag4, which contained some of the same sequence elements as 20a-frag3, we were able to identify correlations for G507 with U508 and with U526, suggesting that G506 is likely unpaired (**Fig. S12**). Interestingly, in this spectrum, we also note the presence of four resonances consistent with non-canonical base pairs that may form in the apical loop, *vide infra*.

### The tertiary structure of FL pre-miR-20a

We next determined the three-dimensional structure of FL pre-miR-20a (**Fig. 2, Table S2**). FL pre-miR-20a adopts a largely elongated, imperfectly paired helical structure, with a dynamic 11-nt apical loop (**Fig. 2A,B**). Sequential aromatic-aromatic and aromatic-anomeric NOEs define the stacking geometry of A547 between U546 and C548 (**Fig. S13**). Additionally, we observed a strong cross-strand NOE between A547.H2 and A486.H1ʹ, which further supports the stacking of A547 in an A-helical geometry (**Fig. S13**). NOEs connecting C485 to the neighboring residues were somewhat sparse, potentially due to spectral overlap. We note that we did not observe an expected NOE between A486.H8 and C485.H1ʹ even in labeled spectra where overlap did not pose a problem. However, we observed a non-canonical NOE between C485.H6 and A486.H1ʹ (**Fig. S13**) which distorts the helical structure, drawing C485 closer to A486.

**Figure 2.**
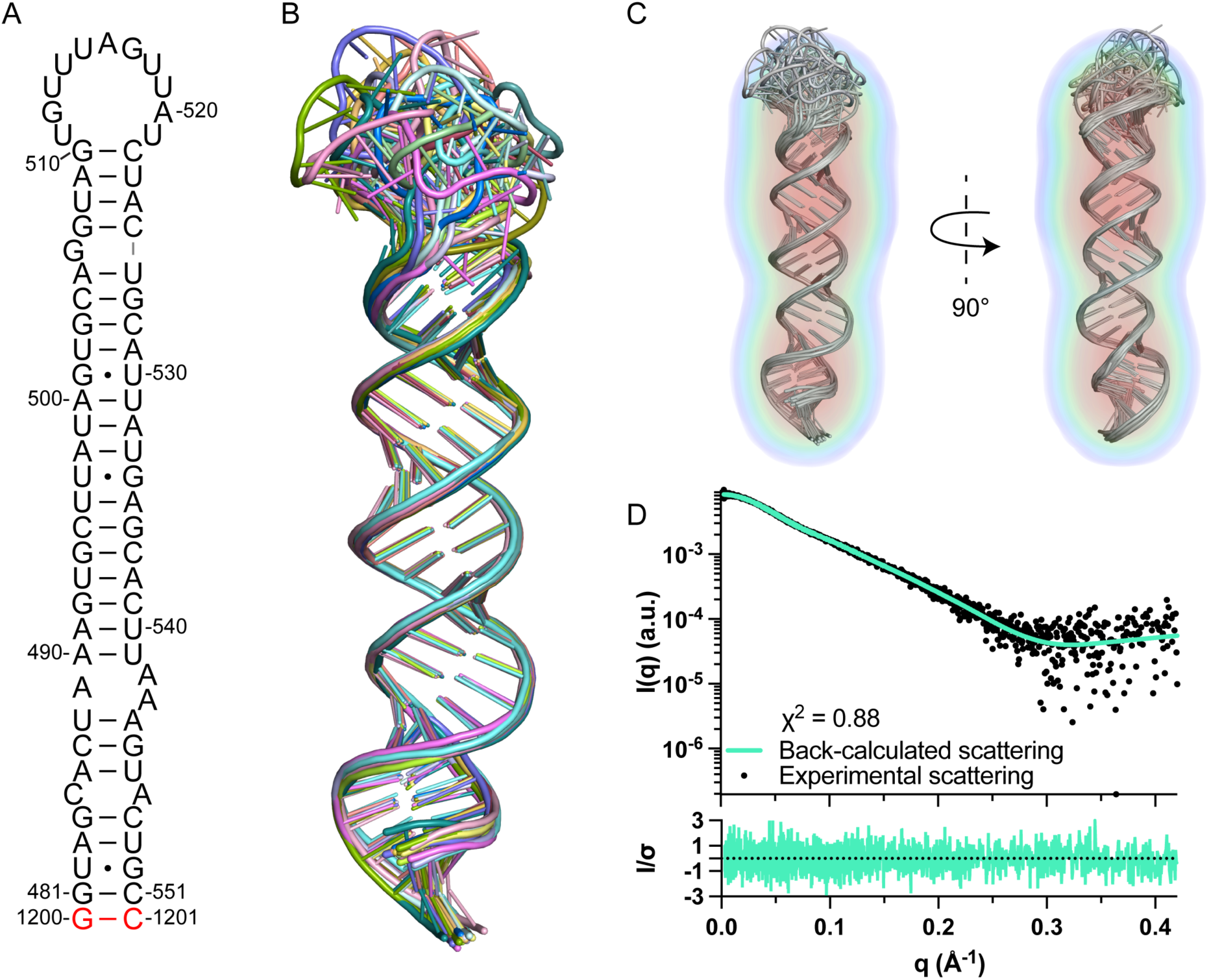
Tertiary structure of FL pre-miR-20a. (A) Secondary structure of FL pre-miR-20a. (B) Ensemble of 20 lowest energy structures. (C) Structure ensemble fit to the SAXS electron density map. (D) The back-calculated scattering curve fits the experimental scattering data (top). Normalized residuals (bottom).

Strong sequential and cross-strand NOEs indicate that A489, A542, and A543 are stacked within a 1×2 internal loop (**Fig. S13**). There were strong aromatic-aromatic NOEs for residues surrounding the G bulge (G506), however, we observed an NOE between A505.H2 and G507.H1ʹ (**Fig. S13,S14**), suggesting that G506 is bulged out which is consistent with our imino analysis of the isolated 20a-frag4 construct (**Fig. S12**).

As an orthogonal method of global structure validation, we used small angle X-ray scattering (SAXS). The refined structure of FL pre-miR-20a was in good agreement with *ab initio* molecular reconstruction derived from SAXS data (**Fig. 2C,D, Table S3**). Kratky analysis suggested that FL pre-miR-20a was well folded, consistent with the derived hairpin structure (**Fig. S15**). The pair-distance distribution function [P(r)] curve revealed the maximum distance (D_max_) to be 112 Å, which matched the overall length of FL pre-miR-20a and the most probable pair distance was ∼ 20 Å, congruent with the average diameter of an A-helical duplex (**Fig. S15**).

### Multiple minor conformations exist in the apical loop of pre-miR-20a

In our analysis of the deuterium-edited ^1^H-^1^H NOESY spectra for FL pre-miR-20a, we noted the presence of strong NOEs between A520.H2 and several cross-strand residues in the ^1^H-^1^H NOESY spectrum, suggesting that there may be additional structure in this region (**Fig. S16**). Interestingly, these NOEs are consistent with two alternative apical loop conformations that we observed in the 20a-frag4 imino spectrum (**Fig. 3A, S12**). While we do not have evidence that these alternative conformations are significantly populated at physiological temperatures, their presence is evident upon lowering the temperature (**Fig. S17**). Furthermore, lowering the pH has a dramatic effect on the relative populations of these distinct states. Integration of the auto peaks of G512b and G512c at pH 6.5 revealed that “conformation b” is present at roughly 60% while “conformation c” is present at roughly 40% (**Fig. 3B**). At pH 7.5, the relative population of these conformers shifts to significantly favor “conformation b” (95%) (**Fig. 3C**). We note that these experiments do not allow us to quantify the population of pre-miR-20a in “conformation a” as G512 is unpaired in this state and therefore does not have a visible imino proton resonance. Together, these findings are consistent with the formation of multiple alternative conformations within the apical loop of pre-miR-20a, which may affect Dicer-TRBP processing to generate mature miR-20a.

**Figure 3.**
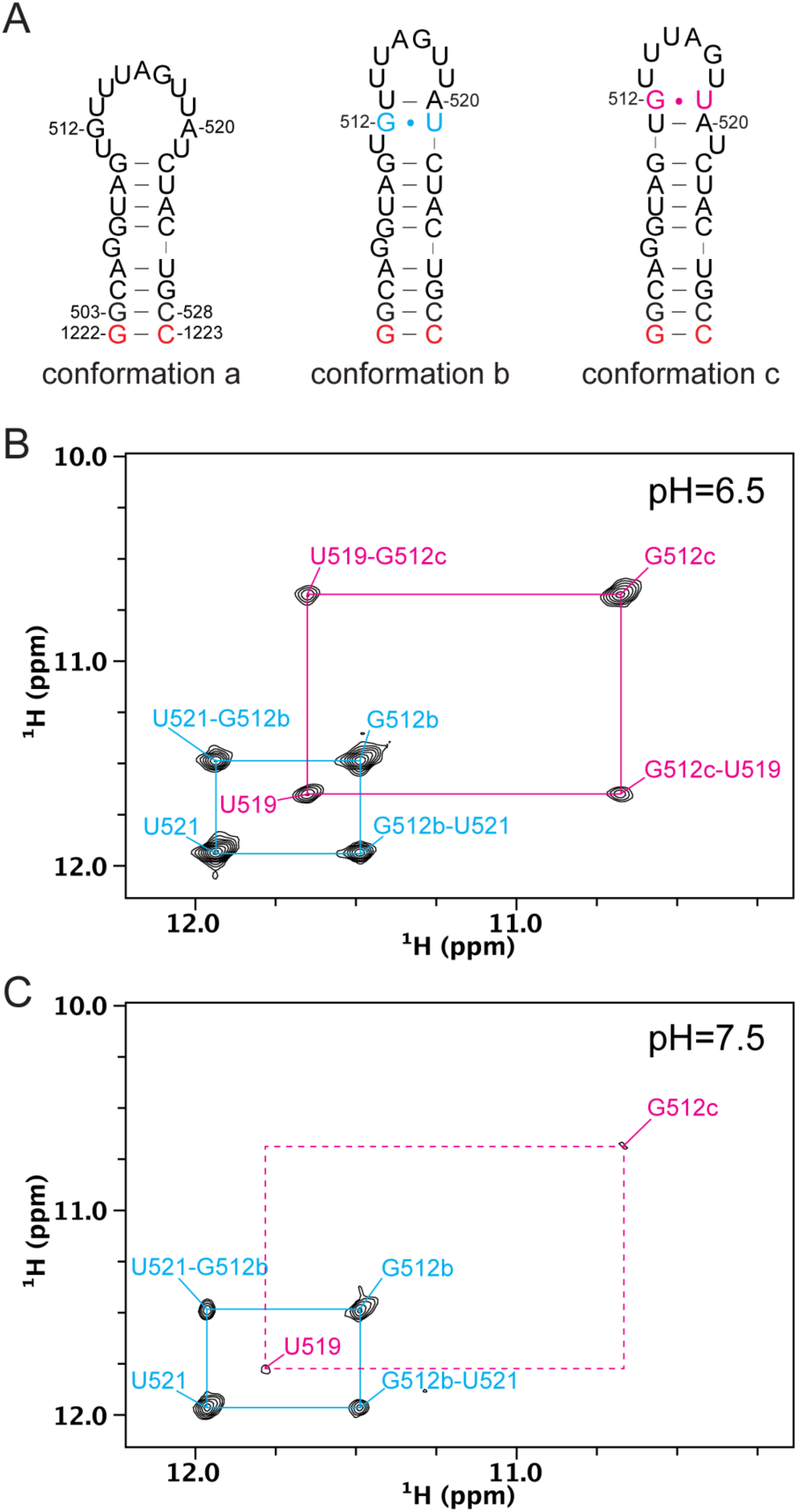
Evidence of alternative conformations within the pre-miR-20a apical loop. (A) Secondary structures depicting three distinct conformations in the apical loop of pre-miR-20a, whose populations are affected by solution conditions. (B,C) Region of the ^1^H-^1^H imino NOESY collected on 20a-frag4 at pH 6.5 (B) and at pH 7.5 (C).

### Apical loop conformations of pre-miR-20a are functionally relevant to Dicer-TRBP processing

The structure of a pre-miRNA apical loop can impact Dicer processing by altering interactions between the upper stem loop of the pre-miRNA and the RNase III and helicase domains of Dicer.^15,28,53^ To investigate the functional relevance of the alternative structures we identified within the pre-miR-20a apical loop, we engineered several mutations to pre-miR-20a intended to stabilize these alternative apical loop structures. (**Fig. 4A**).

**Figure 4.**
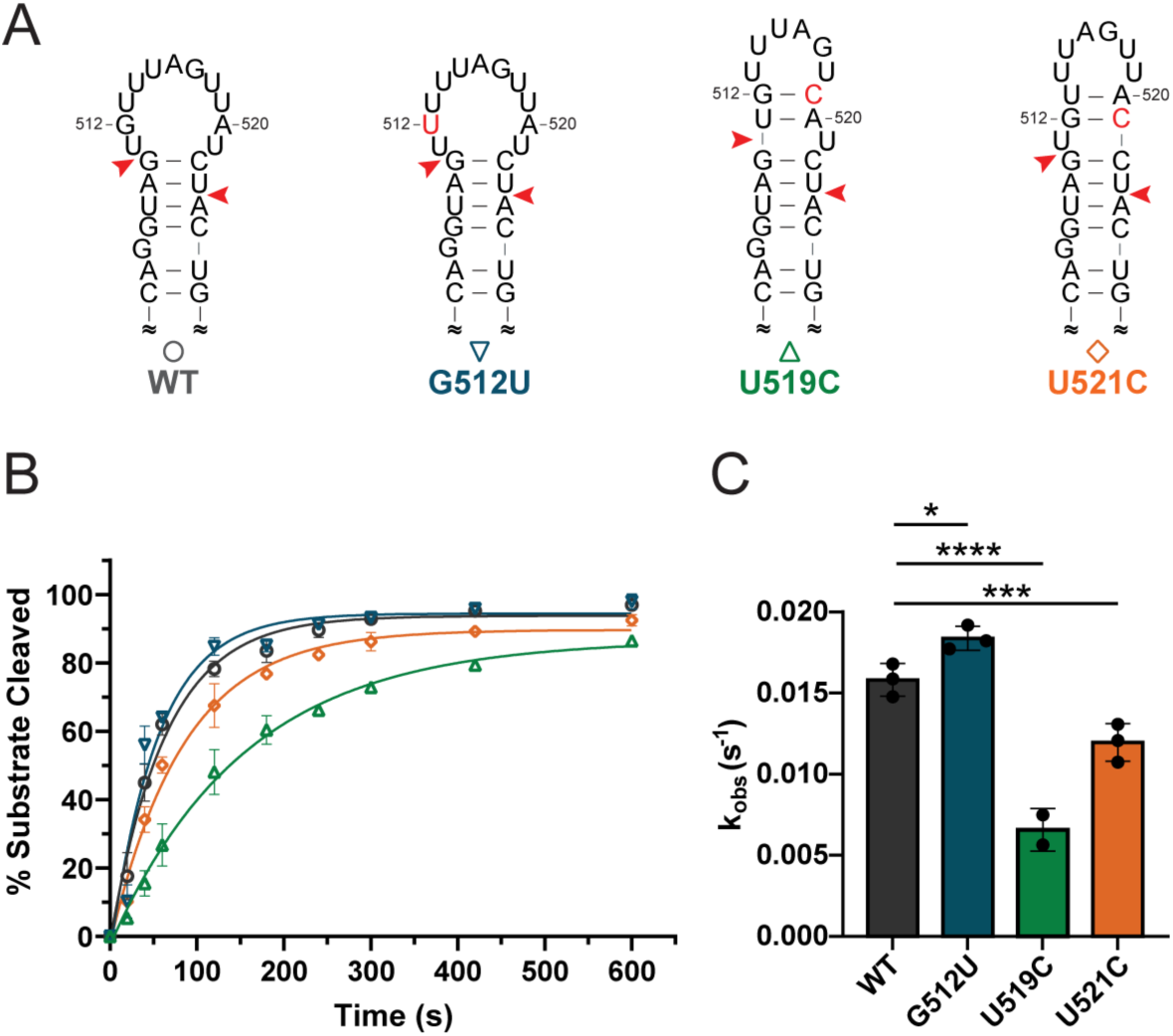
Apical loop conformers affect Dicer-TRBP processing *in vitro*. (A) Predicted secondary structures of WT and mutant pre-miR-20a constructs designed to stabilize distinct apical loop conformations of pre-miR-20a. Nucleotides in red represent mutations, red arrows denote Dicer cleavage sites. (B) Time course of Dicer-TRBP cleavage of pre-miR-20a RNAs. (C) Observed rates (kobs) of Dicer-TRBP cleavage. Data represents mean ± s.d. of 2-3 independent replicates. ****p < 0.0001, ***p < 0.001, *p < 0.05.

We subjected the mutated pre-miR-20a constructs to *in vitro* Dicer-TRBP processing assays to determine the effects of the alternative apical loop structures on processing. We found that the wild-type (WT) pre-miR-20a is processed at a rate of 0.01581 s^-1^ and is near fully processed within 10 minutes (**Fig. 4B,C, Table S10**). The U519C construct, designed to stabilize the apical loop into “conformation c” exhibited a 2.8-fold reduction in Dicer-TRBP processing rate relative to WT (**Fig. 4B,C, Table S10**). Comparatively, the mutation at U521 (U521C) to promote “conformation b” exhibited a more modest 1.3-fold reduction in Dicer-TRBP processing rate relative to WT (**Fig. 4B,C, Table S10**). These findings are consistent with previous studies which demonstrated that apical loops that are smaller than ∼11 nt exhibited reduced Dicer processing.^25,27^ Interestingly, the G512U mutant, designed to promote the formation of “conformation a” exhibited an observed rate constant that was 1.2-fold faster than WT (**Fig. 4B,C, Table S10**). These findings suggest that sampling of alternative apical loop conformations within pre-miR-20a results in functional differences in enzyme processing at physiological temperature and pH, despite their apparent low population.

### A single nucleotide bulge at position 506 is critical for efficient processing of pre-miR-20a

Our 20a-frag3 imino proton data, collected at low temperature, was not consistent with G506 or G507 participating in stable base pairing (**Fig. S11**). Conversely, imino proton data collected on 20a-frag4, which contained the native apical loop structure, was consistent with a bulged G506 and a G507-C525 base pair (**Fig. S12**). At physiological temperature, our NOE data for FL pre-miR-20a support a model in which A505 and G507 are stacked in the A helical structure, but not directly stacked on each other (**Fig. 5A**). G506 is unpaired and protrudes into the minor groove yet is still positioned between A505 and G507. Given the proximity of G506 and G507 to the 5′-dicing site, we next examined whether the base-pairing state of G506 and G507 may impact Dicer-TRBP processing. We generated G506C and G507C mutants of pre-miR-20a to force a single nucleotide bulge at position 506 or 507, respectively (**Fig. 5B**), and subjected them to *in vitro* Dicer-TRBP processing. Pre-miR-20a G506C exhibited WT-like rates of processing (**Fig. 5C,D, Table S10**). However, pre-miR-20a G507C exhibited a 1.8-fold reduction in processing rate relative to WT (**Fig. 5C,D, Table S10**). These results match our 20a-frag4 data and suggest that G507 is engaged in a base-pair with C525, and that the bulge at position 506 is critical for optimal pre-miR-20a processing.

**Figure 5.**
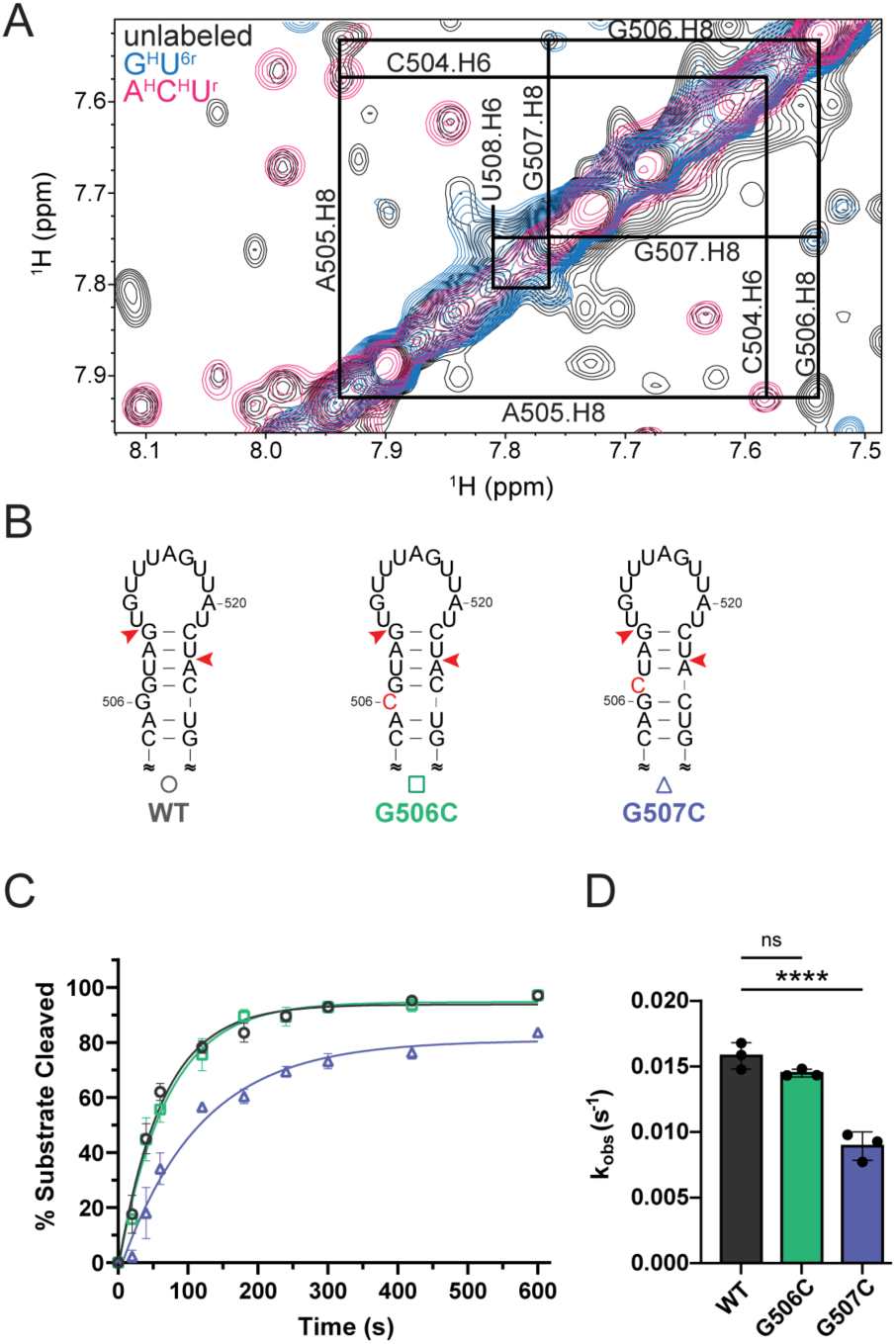
Structure proximal to the dicing site affects Dicer-TRBP processing *in vitro*. (A) Overlay of ^1^H-^1^H NOESY spectra of unlabeled (black), G^H^U^6r–^labeled (blue), and A^H^C^H^U^r^-labeled (pink) FL pre-miR-20a. Sequential aromatic-aromatic NOEs near the dicing site are consistent with stacking. (B) Predicted secondary structures of WT and mutant pre-miR-20a constructs designed to alter the structure near the dicing site. Nucleotides in red represent mutations, red arrows denote Dicer cleavage sites. (C) Dicer-TRBP processing of WT and mutant RNAs as a function of time. (D) Observed rates (kobs) of Dicer-TRBP cleavage for mutant and WT RNAs. Data represents mean ± s.d. of 3 independent replicates. ****p < 0.0001, ***p < 0.001, *p < 0.05.

### A disease-related mutation in pre-miR-20a results in reduced processing efficiency

MiRNAs play important roles in post-transcriptional gene regulation, and the structure of the pre-miRNA, specifically near the dicing site, can greatly influence Dicer processing of the RNA.^30,54,55,25,56^ Using the miRNASNP-v3 database,^57^ we noted a disease related variant of pre-miR-20a associated with colorectal carcinoma (COSN26984469) which contained a U508C mutation near the dicing site. The predicted secondary structure of this mutation shows generation of a large internal loop at the 3ʹ dicing site (**Fig. 6A**) which we predicted would result in an impairment of Dicer-TRBP processing. We find that, indeed, a U508C mutation does reduce Dicer-TRBP processing rate by 3.3-fold compared to WT (**Fig. 6B,C**).

**Figure 6.**
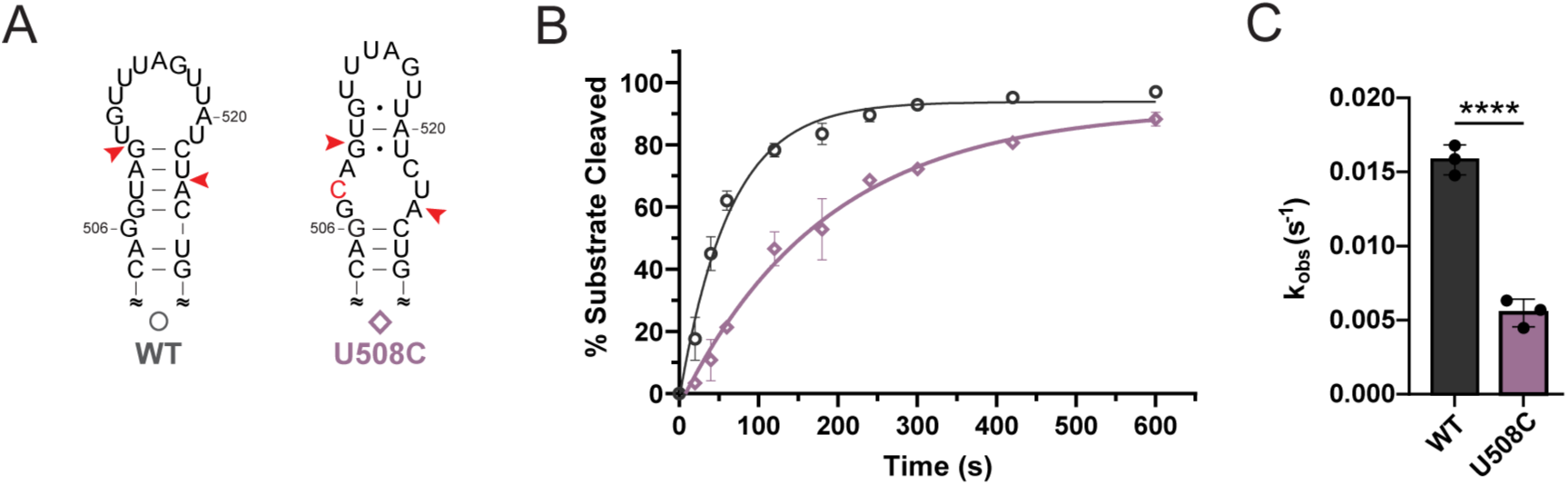
A disease-relevant SNP dramatically reduces Dicer-TRBP processing rate of pre-miR-20a. (A) The pre-miR-20a U508C mutation is predicted to significantly alter the RNA structure near the dicing site. The single point mutation is indicated as a red nucleotide and red arrows denote Dicer cleavage sites. (B) Dicer-TRBP processing time course for WT and U508C pre-miR-20a RNAs. (C) Observed Dicer-TRBP cleavage rates (kobs). Data represents mean ± s.d. of 3 independent replicates, ****p < 0.0001.

## Discussion

RNA dynamics and secondary structure rearrangement are essential to a wide range of cellular processes including protein recruitment and cellular function.^30,58,59^ Regulation of both Drosha and Dicer processing is essential to ensure proper levels of miRNAs are produced. This regulation can be mediated by RNA-protein interactions,^20,60,61^ RNA modification,^24,62–64^ and, as highlighted by this work, conformational rearrangements within pre-miRNAs themselves. Consistent with a growing body of research, we find dynamic interconversion of structures in the apical loop region of pre-miR-20a can tune Dicer-TRBP processing.

Despite their low populations, these alternative structures, which constrict the size of the apical loop to a less optimal Dicer-TRBP substrate, carry functional consequences on *in vitro* enzymatic processing of pre-miR-20a. While we note that removing the ability of pre-miR-20a to adopt these alternative structures through mutation results in a modest increase in processing rate *in vitro*, the effects of such alternative conformations have not been tested *in vivo*. As Dicer-TRBP is responsible for cleaving multiple pre-miRNA species in cells, the cellular environment is inevitably more competitive than our *in vitro* processing assays, and as such small changes in processing *in vitro* could result in more dramatic shifts in *in vivo* processing. Additionally, pre-miR-20a exhibits a comparatively high rate of Dicer-TRBP processing,^16,25,30,31^ and as such increases to the processing rate may not be discernable in our experiments.

An additional hallmark of the pre-miR-20a structure is the presence of a single-nucleotide G bulge proximal to the dicing site. We showed the importance of the positioning of this G bulge for efficient Dicer-TRBP processing and predict this unpaired G may regulate interactions with Dicer-TRBP, as the double-stranded RNA-binding domain (dsRBD) and RNaseIII domain of Dicer is known to interact with this region of pre-miRNAs.^53,65,66^ Additionally, the stability of the base paired region just below the apical loop has been shown to control Dicer processing in pre-miR-31.^25^ An enforced G507 bulge would slightly destabilize the helical region spanning the dicing sites and we found that this destabilized mutant resulted in reduced processing. The importance of the structure and stability of the helical region spanning the dicing sites is particularly clear when we examined the processing of a disease-relevant variant of pre-miR-20a. This single point mutation is predicted to introduce a 3×3 internal loop and disrupt the helical structure at the 3ʹ dicing site. *In vitro*, this mutation was deleterious to Dicer-TRBP processing rate, which is particularly intriguing as higher expression of miR-20a is generally associated with colorectal cancer,^67–70^ although also with better prognosis.^69,70^

## Methods

### Preparation of DNA Templates

Template strand DNAs, containing a T7 RNA promoter and corresponding sequences for the RNA oligonucleotides, were purchased from Integrated DNA Technologies (**Table S4**). Partially-double stranded DNA templates for *in vitro* transcription were created by annealing the template strand DNA with a DNA corresponding to the T7 promoter sequence (5ʹ-TAATACGACTCACTATA-3ʹ). Briefly, the templates were prepared by mixing the desired template strand DNA (40 μL, 200 μM) with the T7 promoter (20 μL, 600 μM), boiling for 3 min, and then slowly cooling to room temperature. The annealed template was diluted with water prior to use to produce the partially double-stranded DNA templates at a final concentration approximately 8 μM.

The double-stranded template corresponding to FL pre-miR-20a (for NMR and SAXS studies) was generated by overlap-extension (OE) PCR using EconoTaq PLUS 2x master mix (Lucigen) with primers listed in **Table S5**.

The native pre-miR-20a template (for processing studies) was generated by OE PCR using EconoTaq PLUS 2x Master Mix (Lucigen) with primers listed in **Table S6**. To ensure homogeneous 5′ and 3′ ends of the RNA, a double ribozyme construct^71^ was generated which contained a hammerhead (HH) ribozyme 5ʹ to the native pre-miR-20a and a hepatitis delta virus (HDV) ribozyme at the 3ʹ end. The PCR template was digested with EcoRI and BamHI restriction enzymes and ligated into the pUC-19 plasmid. All subsequent mutations and deletions were achieved via site-directed mutagenesis (New England Biolabs Q5 site-directed mutagenesis kit) of the HH-pre-miR-20a-HDV plasmid with primers listed in **Table S7**. Templates prepared from plasmids were amplified with EconoTaq PLUS 2x Master Mix (Lucigen) using primers UNIV-pUC19_E105 and HDV-AMP-R (**Table S8**). Plasmid identity was verified by Sanger sequencing (Eurofins Genomics) using the universal M13REV sequencing primer.

### RNA Preparation

RNAs were made using *in vitro* transcription in a 1 x transcription buffer [40 mM Tris base, 5 mM dithiothreitol (DTT), 1 mM spermidine and 0.01% Triton-X (pH 8.5)] with addition of 30-40 ng/μL DNA template, 10-20 mM magnesium chloride (MgCl_2_), 3-6 mM ribonucleoside triphosphates (NTPs), 0.2 unit/mL yeast inorganic pyrophosphatase (New England Biolabs), approximately 15 μM T7 RNA polymerase made in-house, and 10-20% (v/v) dimethyl sulfoxide (DMSO). Reaction mixtures were incubated at 37 °C for 3-4 h, with shaking at 75 rpm, and then quenched using a solution of 7 M urea and 0.5 M ethylenediaminetetraacetic acid (EDTA), pH 8.5. Reactions were boiled for 3 min and then snap-cooled in ice water for 3 min. The transcription mixture was loaded onto preparative-scale 10% or 16% denaturing polyacrylamide gels for purification, depending on the RNA size. RNAs were visualized by UV shadowing and gel slices with RNA were excised. Gels slices were placed into an EluTrap electroelution device (The Gel Company) in 1x TBE buffer. RNA was eluted from the gel at a constant voltage (120 V) for ∼ 24 h. The eluted RNA was spin concentrated, washed with 2 M high-purity sodium chloride (NaCl), and exchanged into water using Amicon Ultra-4 Centrifuge Filter Units (Millipore, Sigma). RNA purity was confirmed on 10% or 16% analytical denaturing gels. RNA concentration was quantified via UV-Vis absorbance. Sequences for all RNAs is provided in **Table S9**.

### Isotopic Labeling of RNAs for NMR

Isotopically labeled RNAs were produced as described above by replacing the rNTP mixture with rNTPs of appropriate isotope labeling. ^15^N rNTPs were obtained from Cambridge Isotope Laboratories (CIL, Andover, MA). The partially- and per-deuterated rNTPs were obtained from Cambridge Isotope Laboratories (CIL, Andover, MA) or prepared in-house, as described below. Deuteration of the C8 position of fully protiated rGTP and rATP (to make G^r^ and A^2r^ rNTPs, respectively) was achieved by incubation with triethylamine (TEA, 5 equiv.) in D_2_O (99.8% deuteration; CIL) for either 24 h (rGTP) or 5 days (rATP). TEA was subsequently removed by lyophilization.

### NMR Experiments

All the RNAs were boiled, then snap-cooled in ultra-pure water. NMR samples for 20a-frag1, 20a-frag2, 20a-frag3 and 20a-frag4, and FL pre-miR-20a were prepared in buffer containing 50 mM K-phosphate buffer, pH 7.5, 1 mM MgCl_2_ in either 99.8% D_2_O (CIL) or 10% D_2_O/90% H_2_O. RNAs were prepared at a concentration of 400-800 μM. NMR spectra were collected on 600 and 800 MHz Bruker AVANCE NEO spectrometers equipped with a 5 mm three channel inverse (TCI) cryogenic probe (University of Michigan BioNMR Core). NMR data were processed with NMRFx^72^ and analyzed with NMRViewJ.^73^ ^1^H chemical shifts were referenced to water and ^13^C chemical shifts were indirectly referenced from the ^1^H chemical shift.^74^

The signals of nonexchangeable protons of 20a oligos were assigned based on analysis of 2D ^1^H-^1^H NOESY (τ_m_ = 300 ms), 2D ^1^H-^1^H TOCSY (τ_m_ = 80 ms), and 2D ^1^H-^13^C HMQC spectra collected at 37 °C. Nonexchangeable proton assignments of FL pre-miR-20a were obtained from 2D NOESY data (τ_m_ ranging from 150-400 ms, 37 °C) recorded on fully protiated, A^2r^G^r^C^r^-, A^H^C^H^U^r^-, and G^H^U^6r–^labeled pre-miR-20a (superscripts denote sites of protiation on a given nucleoside, all other sites deuterated). A best-selective long-range HNN-COSY^45^ was recorded on 750 μM ^15^N A,U-labeled pre-miR-20a in 10% D_2_O/90% H_2_O, 50 mM K-phosphate buffer, pH 7.5, 1 mM MgCl_2_.

### Structure Calculation

CYANA was used to generate 800 initial structures via simulated annealing molecular dynamics calculations over 128,000 steps. Upper limits for the NOE distance restraints were generally set at 5.0 Å for weak, 3.3 Å for medium, and 2.7 Å for strong signals, based on peak intensity. For very weak signals, 6.0 Å upper limit restraints were used, including for sequential H1′–H1′ NOEs and intraresidue H5–H1′ NOEs. Standard torsion angle restraints were included for regions with A-helical geometry, allowing for ±25° deviations from ideality (ζ=−73°, α=−62°, β=180°, γ=48°, δ=83°, ɛ=−152°). Torsion angles for mismatches were further relaxed to allow for ±75° deviation from ideality. Standard hydrogen bonding, ribose, and planarity restraints were included. Cross-strand phosphate–phosphate distance restraints were employed for regions of the structure that adopted A-form helical geometry to prevent the generation of structures with collapsed major grooves.

The top 20 CYANA-derived structures were then subjected to molecular dynamics simulations and energy minimization with AMBER.^75^ Only upper limit NOE, hydrogen bond, and chirality restraints were used. Backbone torsion angle and phosphate-phosphate restraints were excluded during AMBER refinement. Calculations were performed using the generalized Born force fields. NMR restraints and structure statistics are presented in **Table S2**.

### SEC-MALS-SAXS Data Acquisition and Analysis

Full details of SAXS data collection and analysis are presented in **Table S3** according to publication guidelines.^76^ SAXS with in-line size exclusion chromatography (SEC) and multiangle light scattering (MALS) was performed at BioCAT (beamline 18ID at the Advanced Photon Source, Chicago). SAXS buffer contained 50 mM potassium phosphate buffer, pH 7.5, 50 mM NaCl, 1 mM MgCl_2_, and all data were collected at 22 °C. The samples were loaded on a Superdex 75 10/300 Increase column (Cytiva) run by a 1260 Infinity II HPLC (Agilent Technologies) at 0.6 mL/min. The flow passed through (in order) the Agilent UV detector, a MALS detector and a DLS detector (DAWN Helios II, Wyatt Technologies), and an RI detector (Optilab T-rEX, Wyatt). The flow then went through the SAXS flow cell. The flow cell consists of a 1.0 mm ID quartz capillary with ∼20 μm walls. A coflowing buffer sheath is used to separate sample from the capillary walls, helping prevent radiation damage.^77^ Scattering intensity was recorded using an Eiger2 XE 9M (Dectris) detector which was placed 3.663 m from the sample giving us access to a q-range of 0.0029 Å^-1^ to 0.42 Å^-1^. 0.5 s exposures were acquired every 1 s during elution and data was reduced using BioXTAS RAW 2.1.4.^78^ Buffer blanks were created by averaging regions flanking the elution peak and subtracted from exposures selected from the elution peak to create the *I*(*q*) vs *q* curves used for subsequent analyses. Molecular weights and hydrodynamic radii were calculated from the MALS and DLS data respectively using the ASTRA 7 software (Wyatt). Guinier and dimensionless Kratky analyses were carried out in BioXTAS RAW (2.2.1). The molecular mass estimates were obtained from the volume of correlation (*V_c_*). The GNOM package within ATSAS (3.0.3)^79^ was used to determine the pair-distance distribution function [*P*(*r*)] required for molecular reconstruction.^80^ DENSS was used to calculate 3D electron density reconstructions. For DENSS, 20 reconstructions were performed in slow mode using default parameters and averaged. Density reconstructions are contoured as follows: 15σ (red), 10σ (yellow), 7.5σ (green), 5σ (cyan), and 2.5σ (blue). Alignment of the reconstruction to the structure was achieved using the DENSS alignment function in BioXTAS RAW. Reconstructions were visualized using PyMOL 3.1.4.1. Theoretical scattering profiles of the NMR structures were back-calculated using FoXS.^81,82^

### Preparation of Recombinant Human Dicer

Expi293 cells expressing FLAG-Dicer were purchased from the University of Michigan Protein core. Cell pellets were lysed via sonication in 20 mM Tris pH 7.4, 150 mM NaCl, 10% glycerol, 1% NP-40, and 0.5 mM TCEP with EDTA-free protease inhibitor. Following sonication, lysate was rocked for 45 min at 4 °C. Anti-FLAG resin was prepared by washing with 1 x TBS (50 mM Tris pH 7.5, 150 mM NaCl), followed by washing with 0.1 M glycine and again with TBS. Lysates were centrifuged at 20,000×*g* for 45 min, and supernatant added to M2 anti-FLAG resin (500 μL/20 mL lysate) for 4 h at 4 °C. Slurry was centrifuged at 500 x g for 5 min and washed with TBS. Resin-bound Dicer was eluted for ∼12-16 h at 4 °C in elution buffer (1 x TBS + 0.5mM TCEP) and resin washed five times with elution buffer. Elutions were concentrated to ∼400 μL and concentration measured by Bradford assay.

### Expression and Purification of TRBP

The expression and purification of human TRBP was based on previously described procedures.^14,25,83,84^ The pET28a-TRBP was purchased from Addgene (Addgene plasmid #50351).

The pET28a-TRBP plasmid was transformed into *Escherichia coli* Rosetta (DE3) pLysS (Novagen), and cells were grown in LB media until cell density reached OD600 = 0.6. Protein expression was induced by the addition of 0.2 mM IPTG and cells were incubated at 18 °C for 20h. The cells were harvested by centrifugation, resuspend in buffer A (20 mM Tris–HCl pH 8.0, 500 mM NaCl, 25 mM imidazole, and 5 mM β-mercaptoethanol), and lysed by sonication. The lysate was pelleted by centrifugation at 20,000×*g* for 30 min, and the supernatant was loaded onto a nickel affinity column and gradient eluted in buffer B (20 mM Tris–HCl pH 8.0, 500 mM NaCl, 250 mM imidazole buffer, and 5 mM β-mercaptoethanol). The protein elution was collected and 10% polyethyleneimine (PEI) was added dropwise (1.5% vol/ vol) to remove nucleic acid contaminants. The suspension was stirred at 4 °C for 30 min and the supernatant was collected by centrifugation (30 min at 20,000 rpm). Two sequential ammonium sulfate cuts were performed (with centrifugation for 30 min at 20,000 rpm in between) at 20% and 80% saturation. The pellet from the 80% ammonium sulfate cut was resuspended in buffer C (20 mM Tris-HCl pH 8.0, 2 mM DTT) and dialyzed against buffer D (20 mM Tris-HCl pH 8.0, 150 mM NaCl, 2 mM DTT) overnight. The overnight dialyzed sample was concentrated and loaded into HiLoad 16/600 Superdex 200 (Cytiva) equilibrated with buffer D.

### Dicer–TRBP Complex Formation

Purified Dicer and TRBP proteins were mixed at a 1:3 molar ratio in 1 mL and concentrated in an Amicon Ultra-4 filter unit (MilliporeSigma) to 100 µL and loaded into HiLoad 16/600 Superdex 200 column (Cytiva) equilibrated with buffer D. The fractions with complex were pooled and concentrated to ∼913 nM, flash-frozen in liquid nitrogen and stored at -80°C.^32^P Labeling of RNA.

The 5′-end labeling of RNA was performed using 5 pmol of RNA, 1 μL 3000 Ci/mmol γ-^32^P-ATP (Revvity), and 10 U T4 polynucleotide kinase (New England Biolabs) in a final volume of 10 μL. RNA was boiled for 3 min and snap-cooled by placing on ice for another 3 min prior to labeling. The radiolabeled RNA was purified on a G-25 column (Cytiva) according to the manufacturer’s instructions. The concentration of radiolabeled RNA was determined using a standard curve derived from the γ-^32^P-ATP source. Radiolabeled RNAs were diluted to 1 nM, and 1 µL combined with cold-labeled RNA (prepared as previously described using unlabeled ATP) to a final total RNA concentration of 20 nM. RNAs were heated to 95 °C for 3 min followed by snap-cool on ice for 3 min and were stored at -20 °C.

### Dicer–TRBP Processing Assay

Concentrated recombinant human Dicer-TRBP complex was diluted to 200 nM in dicing buffer (24 mM HEPES or 24 mM Bis-Tris, pH 7.5, 100 mM NaCl, 5 mM MgCl2, 4 μM EDTA). Diluted Dicer-TRBP complex (1 µL) was premixed with 80 U RNaseOUT Recombinant Ribonuclease Inhibitor (Thermo Fisher Scientific), 2 µL 5x dicing buffer, and water to final volume of 9 µL. The RNA (1 μL) was added to premixed solution (9 μL) and incubated at 37 °C. The final RNA and enzyme concentrations were 2 nM and 20 nM, respectively. The reaction was quenched by adding 10 μL quench buffer (98% formamide, 20 mM EDTA, trace bromophenol blue and xylene cyanol) at 20, 40, 60, 120, 180, 240, 300, 420, and 600 s, respectively. Samples were run on a 12% denaturing polyacrylamide gel, and the gel was exposed to a phosphor screen for ∼24 h followed by scanning on an Amersham Typhoon Phosphor Imager (GE Healthcare). The gel image was quantified by ImageJ. The Dicer cleavage ratio was calculated as the sum of the intensity of products divided by the sum of the intensity of the products, partially digested products, and remaining substrate. Experiments were performed in triplicate unless indicated. The average and SD of the measurements are reported.

The pre-miRNA cleavage ratio was fitted as function of time to get apparent rate constant (*k*_obs_) for each replicate using Graphpad Prism Software. Data are fitted using Eq. 1:

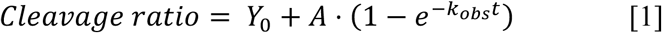

In the fitting parameter, *Y_0_* and *A* are nonregressive fitting parameters are unrestricted when fitting. A one-way ANOVA analysis followed by Dunnett’s multiple comparisons test was used to determine statistical significance between processing rates of mutant RNAs to WT.

### CD Thermal Denaturation

CD thermal denaturation of RNAs was performed on JASCO J1500CD spectrometer with a heating rate of 1 ℃ per min from 5 ℃ to 95 ℃. Data points were collected every 0.5 ℃ with the wavelength set at 260 nm. 20 μM RNA samples were premixed in 50 mM K-phosphate buffer (pH 5.8, 6.5, or 7.5), 1 mM MgCl_2_. The single transition unfolding melting profiles were analyzed using a two-state model using sloping baselines (Eq. 2).^25^

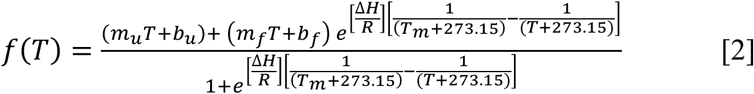

Here, m_u_ and m_f_ are the slopes of the lower (unfolded) and upper (folded) baselines, b_u_ and b_f_ are the y-intercepts of the lower and upper baselines, respectively. ΔH (in kcal/mol) is the enthalpy of folding and T_m_ (in °C) is the melting temperature. Experiments were performed in triplicate. The average, and standard deviation of the measurements are reported.

### Secondary Structure Rendering

All secondary structures were generated using RNACanvas.^85^

### Accession Numbers

Resonance assignments have been deposited in the BMRB (miR-20a_frag1: 52278, frag2: 52279, frag3: 52280, frag4: 52281, and FL pre-miR-20a: 52285). NMR-derived structures of FL pre-miR-20a have been deposited in the PDB (9OBM). Experimental SAXS data of FL pre-miR-20a have been deposited in the Small Angle Scattering Biological Data Bank under accession code SASXXXX.

## Acknowledgements

This work was supported by National Institute of General Medical Sciences of the National Institutes of Health grant R35 GM138279 (to S.C.K.) and Research Corporation for Science Advancement Cottrell Scholar Award 28248 (S.C.K.). Research reported in this publication was supported by the University of Michigan BioNMR Core Facility (U-M BioNMR). U-M BioNMR Core is grateful for support from U-M including the College of Literature, Sciences and Arts, Life Sciences Institute, College of Pharmacy and the Medical School along with the U-M Biosciences Initiative. This research used resources of the Advanced Photon Source, a U.S. Department of Energy (DOE) Office of Science User Facility operated for the DOE Office of Science by Argonne National Laboratory under Contract No. DE-AC02-06CH11357. Research reported in this publication was supported by grant P30 GM138395 from the National Institute of General Medical Sciences of the National Institutes of Health. Use of the Pilatus 3 1M detector was provided by grant 1S10OD018090-01 from NIGMS. The content is solely the responsibility of the authors and does not necessarily reflect the official views of the National Institute of General Medical Sciences or the National Institutes of Health.

We thank Dr. Jesse Hopkins and Dr. Maxwell Watkins at Argonne National Laboratory for assistance with SAXS data collection and analysis. Additionally, we thank Dr. Debashish Sahu and Dr. Minli Xing at the University of Michigan BioNMR Core for their assistance with NMR data collection and instrument maintenance.

