## Supplementary Data for "Structural features within precursor microRNA-20a regulate Dicer-TRBP processing"

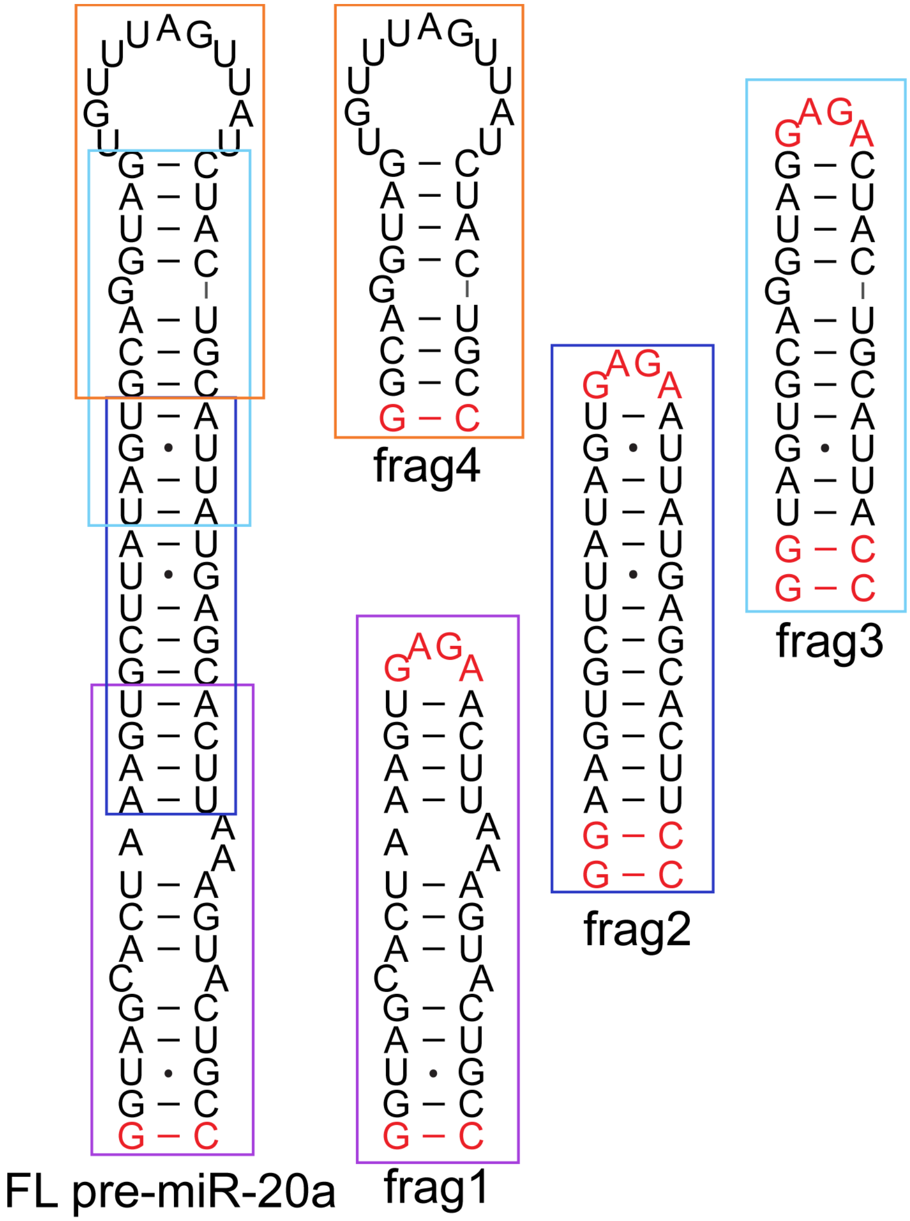

**Figure S1. Oligonucleotide controls for NMR chemical shift assignment of FL pre-miR-20a.** Four oligonucleotide controls were designed to cover the entire FL pre-miR-20a sequence. Non-native tetraloops (red) were included to cap oligos that truncated the apical loop.

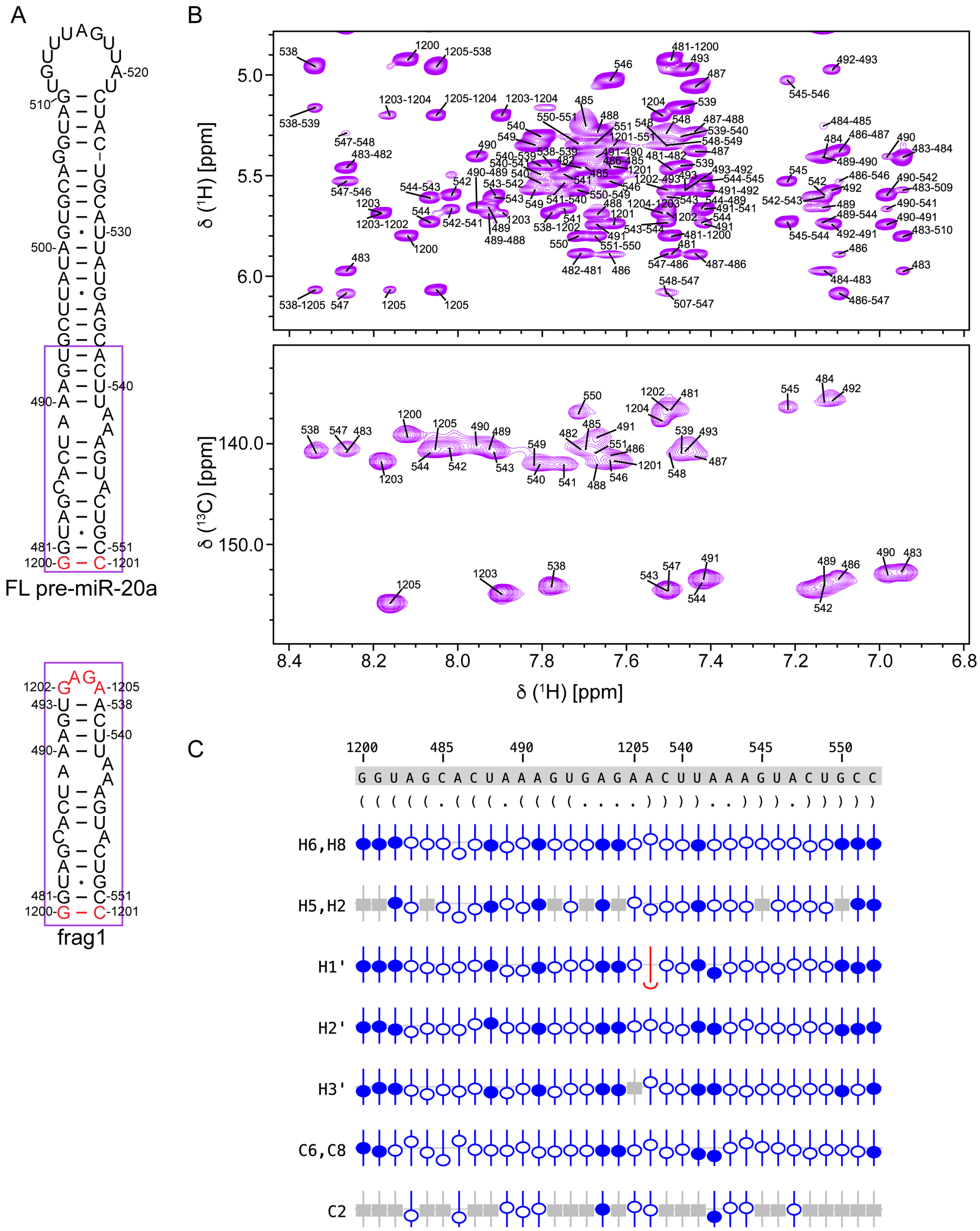

**Figure S2**. **Assigned chemical shifts of frag1.** (A) Secondary structure of FL pre-miR-20a and 20a-frag1. (B) $^{1}$H-$^{1}$H NOESY (top) and $^{1}$H-$^{13}$C HMQC (bottom) of 20a-frag1. (C) Sequence analysis and validation of 20a-frag1 chemical shift assignments. Nucleotide numbering, sequence, and secondary structure in Vienna format for 20a-frag1 RNA. NMRViewJ chemical shift prediction software was used to validate proton (H6/H8, H5/H2, H1′, H2′, H3′) and carbon (C6/C8, C2) assignments. Assigned atoms are represented with blue circles (open and closed), while grey boxes denote atoms that are not present in a given base. Deviation from the predicted chemical shift is shown with the offset from the center. Filled circles indicate that there are chemical shifts for atoms with the same set of attributes in the BMRB. Open circles indicate atoms that have a prediction, but for which no exact matches of the attributes are available in the BMRB.

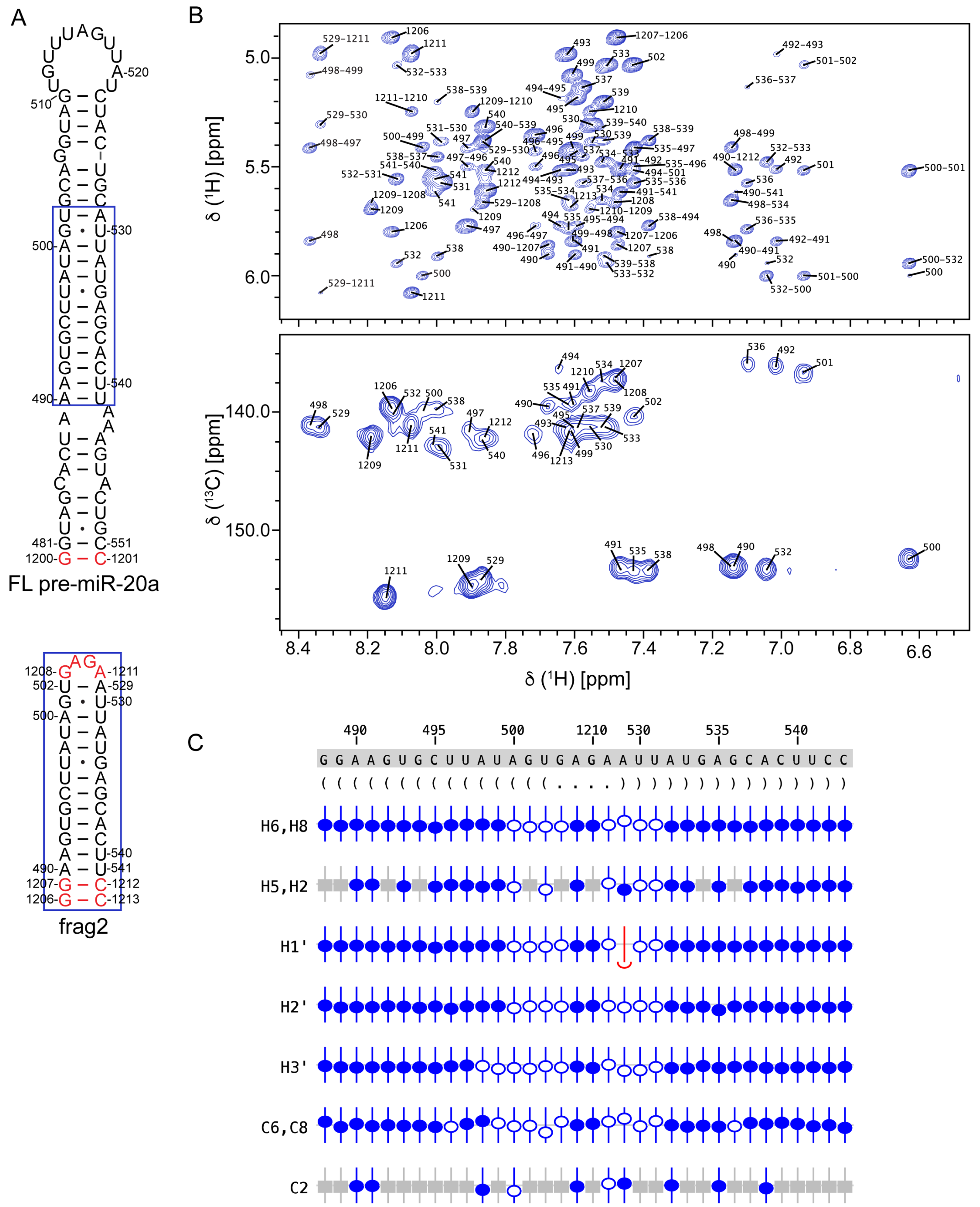

**Figure S3**. **Assigned chemical shifts of frag2.** (A) Secondary structure of FL pre-miR-20a and 20a-frag2. (B) $^{1}$H-$^{1}$H NOESY (top) and $^{1}$H-$^{13}$C HMQC (bottom) of 20a-frag2. (C) Sequence analysis and validation of 20a-frag2 chemical shift assignments. Nucleotide numbering, sequence, and secondary structure in Vienna format for 20a-frag2 RNA. NMRViewJ chemical shift prediction software was used to validate proton (H6/H8, H5/H2, H1′, H2′, H3′) and carbon (C6/C8, C2) assignments. Assigned atoms are represented with blue circles (open and closed), while grey boxes denote atoms that are not present in a given base. Deviation from the predicted chemical shift is shown with the offset from the center. Filled circles indicate that there are chemical shifts for atoms with the same set of attributes in the BMRB. Open circles indicate atoms that have a prediction, but for which no exact matches of the attributes are available in the BMRB.

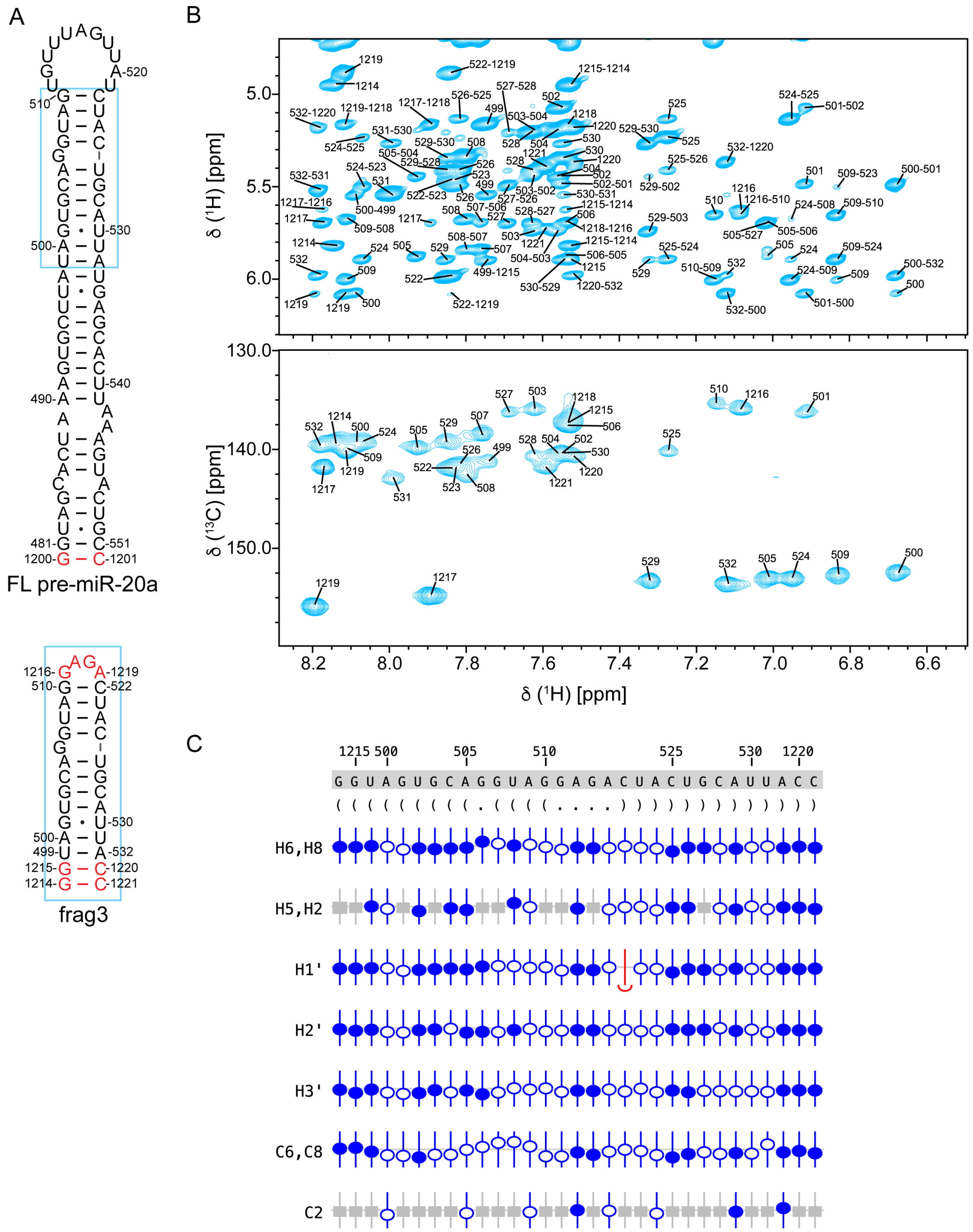

**Figure S4**. **Assigned chemical shifts of frag3.** (A) Secondary structure of FL pre-miR-20a and 20a-frag3. (B) $^{1}H$-$^{1}H$ NOESY (top) and $^{1}H$-$^{13}C$ HMQC (bottom) of 20a-frag3. (C) Sequence analysis and validation of 20a-frag3 chemical shift assignments. Nucleotide numbering, sequence, and secondary structure in Vienna format for 20a-frag3 RNA. NMRViewJ chemical shift prediction software was used to validate proton (H6/H8, H5/H2, H1′, H2′, H3′) and carbon (C6/C8, C2) assignments. Assigned atoms are represented with blue circles (open and closed), while grey boxes denote atoms that are not present in a given base. Deviation from the predicted chemical shift is shown with the offset from the center. Filled circles indicate that there are chemical shifts for atoms with the same set of attributes in the BMRB. Open circles indicate atoms that have a prediction, but for which no exact matches of the attributes are available in the BMRB.

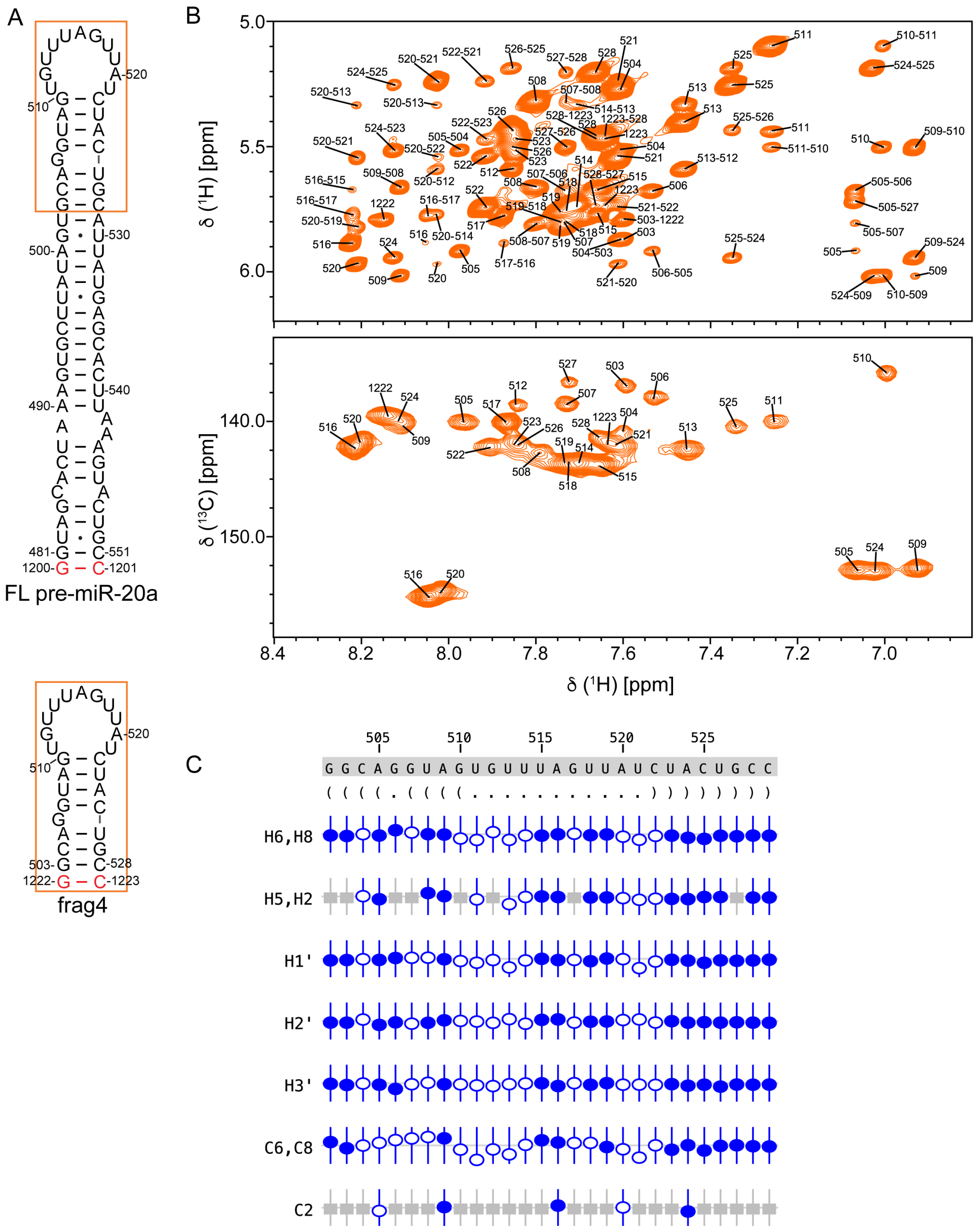

**Figure S5**. **Assigned chemical shifts of frag4.** (A) Secondary structure of FL pre-miR-20a and 20a-frag4. (B) $^{1}$H-$^{1}$H NOESY (top) and $^{1}$H-$^{13}$C HMQC (bottom) of 20a-frag4. (C) Sequence analysis and validation of 20a-frag4 chemical shift assignments. Nucleotide numbering, sequence, and secondary structure in Vienna format for 20a-frag4 RNA. NMRViewJ chemical shift prediction software was used to validate proton (H6/H8, H5/H2, H1′, H2′, H3′) and carbon (C6/C8, C2) assignments. Assigned atoms are represented with blue circles (open and closed), while grey boxes denote atoms that are not present in a given base. Deviation from the predicted chemical shift is shown with the offset from the center. Filled circles indicate that there are chemical shifts for atoms with the same set of attributes in the BMRB. Open circles indicate atoms that have a prediction, but for which no exact matches of the attributes are available in the BMRB.

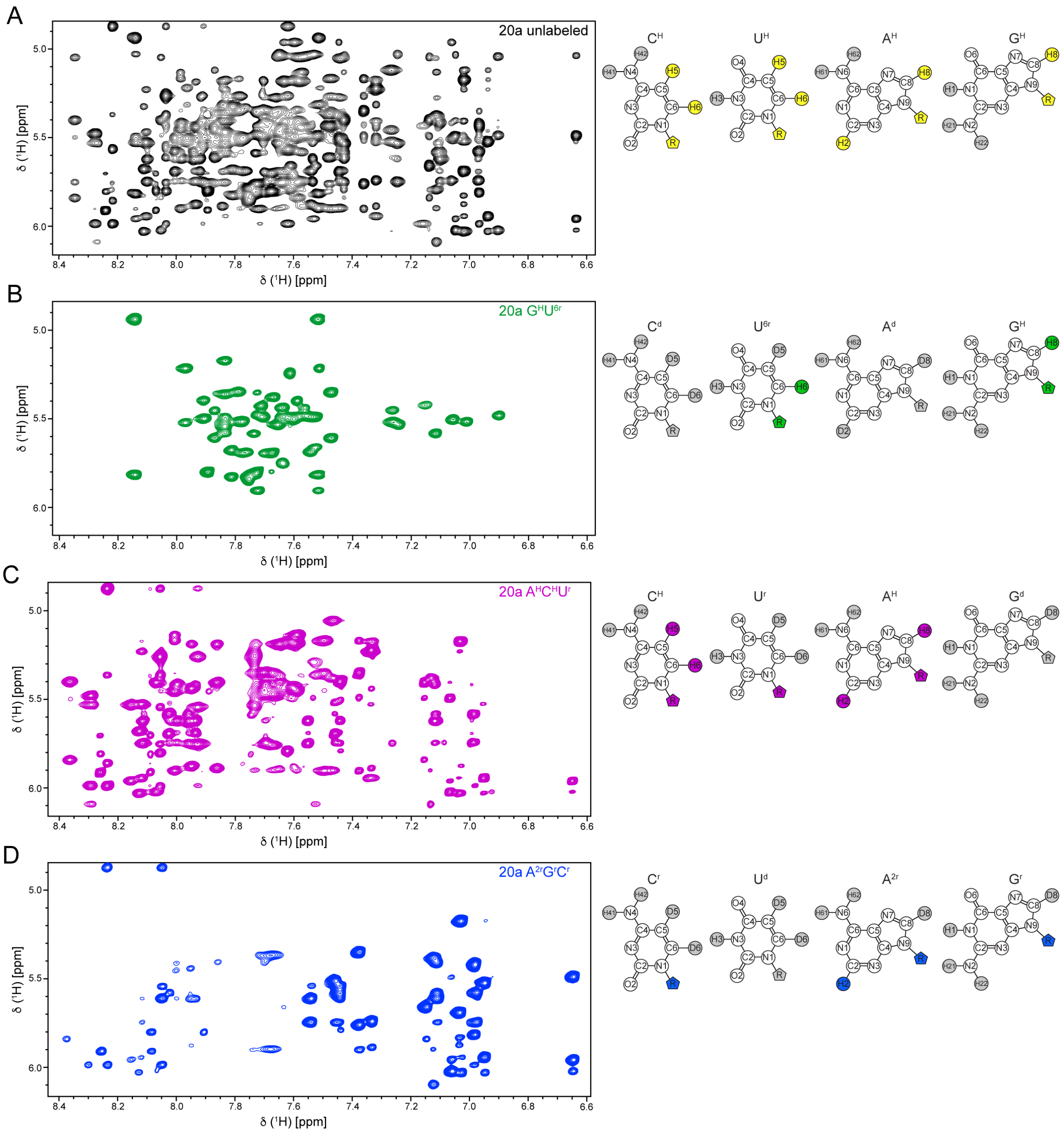

**Figure S6**. **Deuterium labeling improves spectral quality by reducing overlap.** $^{1}H$-$^{1}H$ NOESY spectra of (A) unlabeled (fully protiated), (B) $G^{H}U^{6r}$-labeled, (C) $A^{H}C^{H}U^{r}$-labeled, and (D) $A^{2r}G^{r}C^{r}$-labeled FL pre-miR-20a RNAs. Chemical structures of the four nucleosides are shown to the right of each spectrum. Sites of the selective deuteration and exchangeable protons are shaded gray while non-exchangeable protons are colored according to each spectrum.

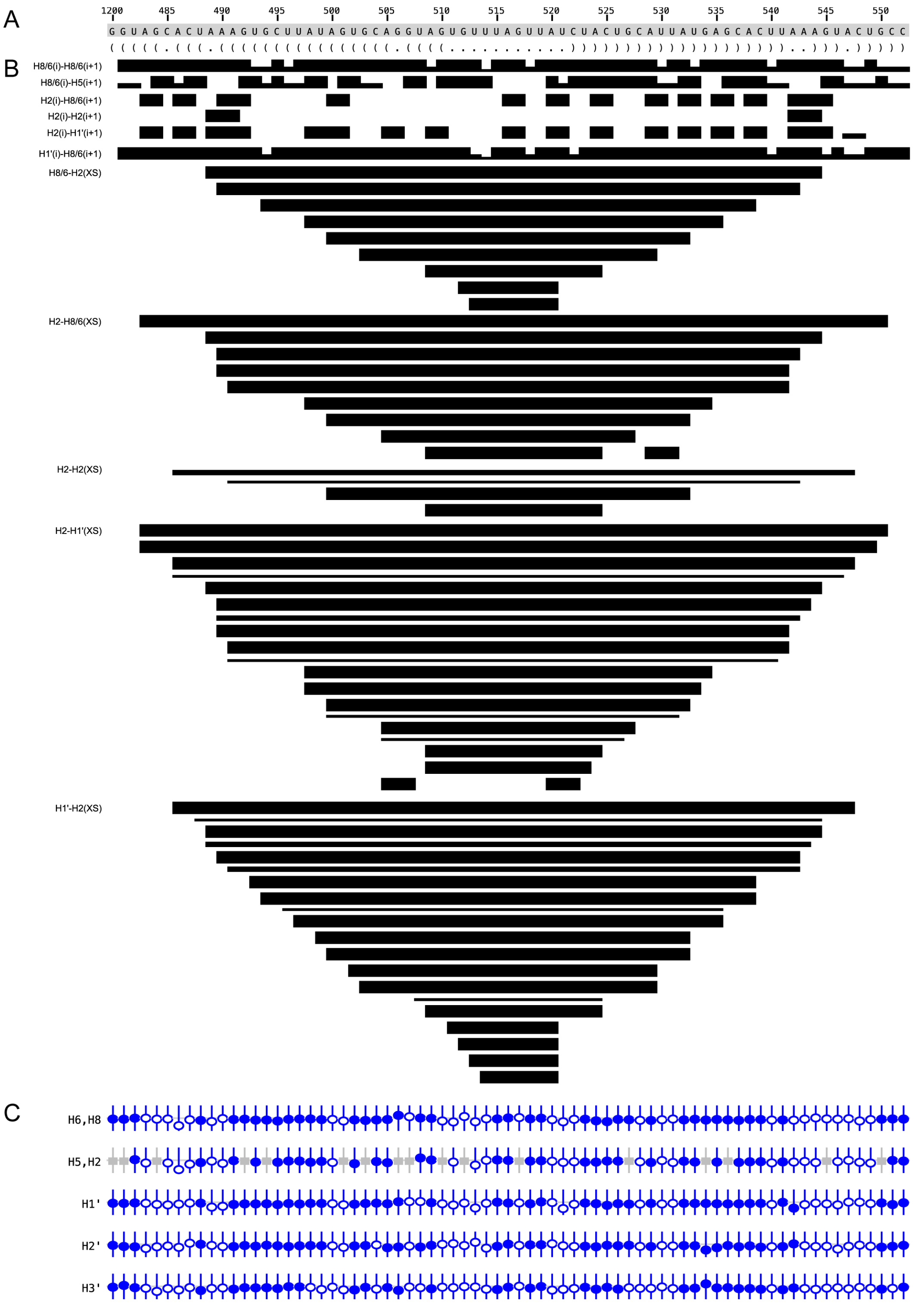

**Figure S7. Summary of the secondary structure, NOE connectivity, and chemical shift assignment validation for FL pre-miR-20a.** (A) Sequence and secondary structure of FL pre-miR-20a. (B) Inter-residue NOE upper limit restraints for specified proton pairs used in structure calculations are denoted with black bars. The thickness of the bar is representative of the strength of the measured NOE. (C) Chemical shift validation from NMRViewJ.

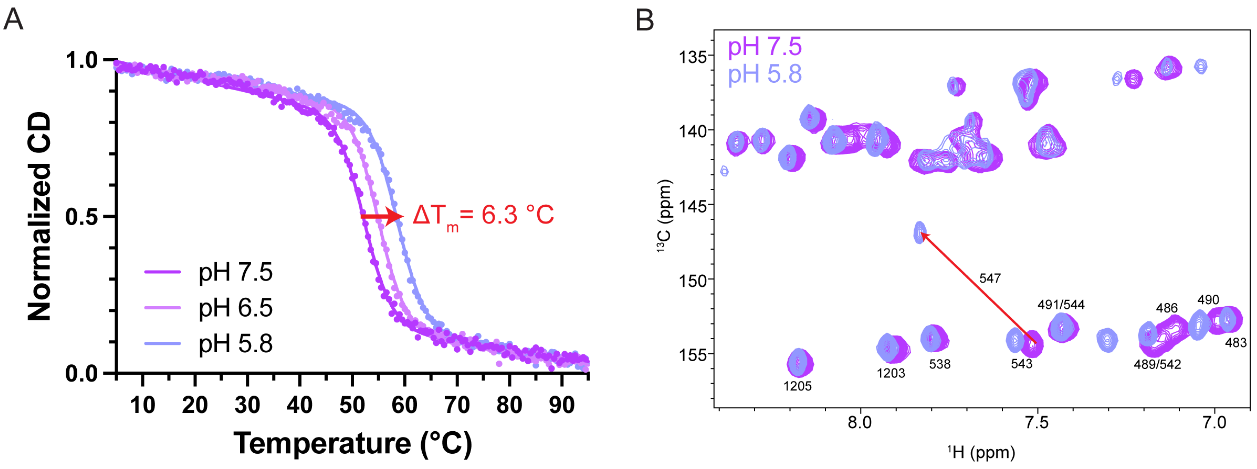

**Figure S8. A547 is protonated at low pH and forms a non-canonical base pair with C485.** (A) Normalized CD-thermal denaturation curves of 20a-frag1 at pH 7.5 (magenta), pH 6.5 (pink), and pH 5.8 (blue). (B) $^{1}H$-$^{13}C$ HMQC overlay of 20a-frag1 at pH 7.5 (magenta) and pH 5.8 (blue). Arrow indicates the significant shift of the A547 C2-H2 peak consistent with protonation at low pH.

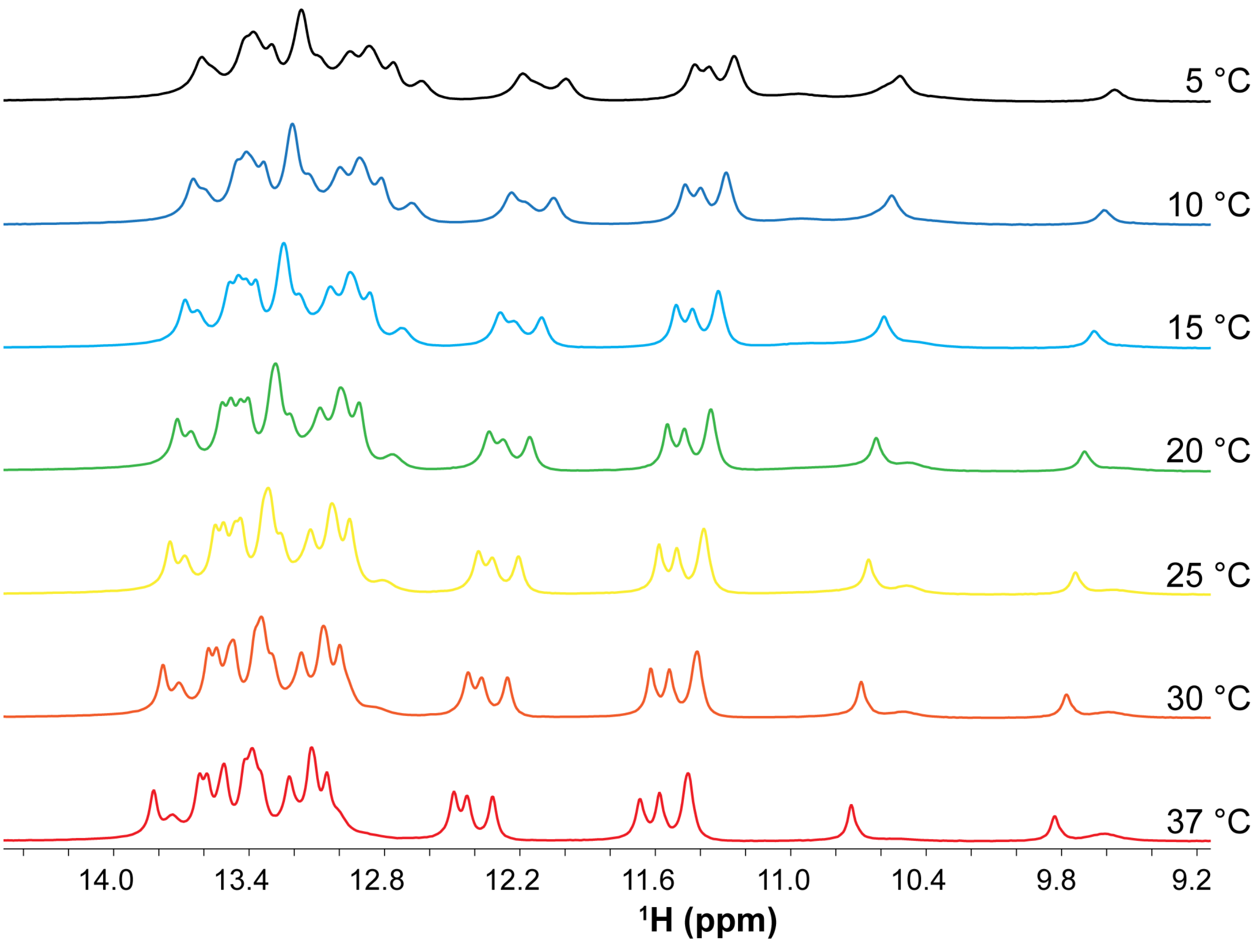

**Figure S9. Imino region of $^1$H spectra of FL pre-miR-20a as a function of temperature.** The NMR spectra were recorded at 0.4 mM RNA concentration, 50 mM K-phosphate buffer, pH 6, 1 mM $MgCl_2$ and 90% $H_2O$/10% $D_2O$ at 600 MHz and at temperatures between 5 °C and 37 °C.

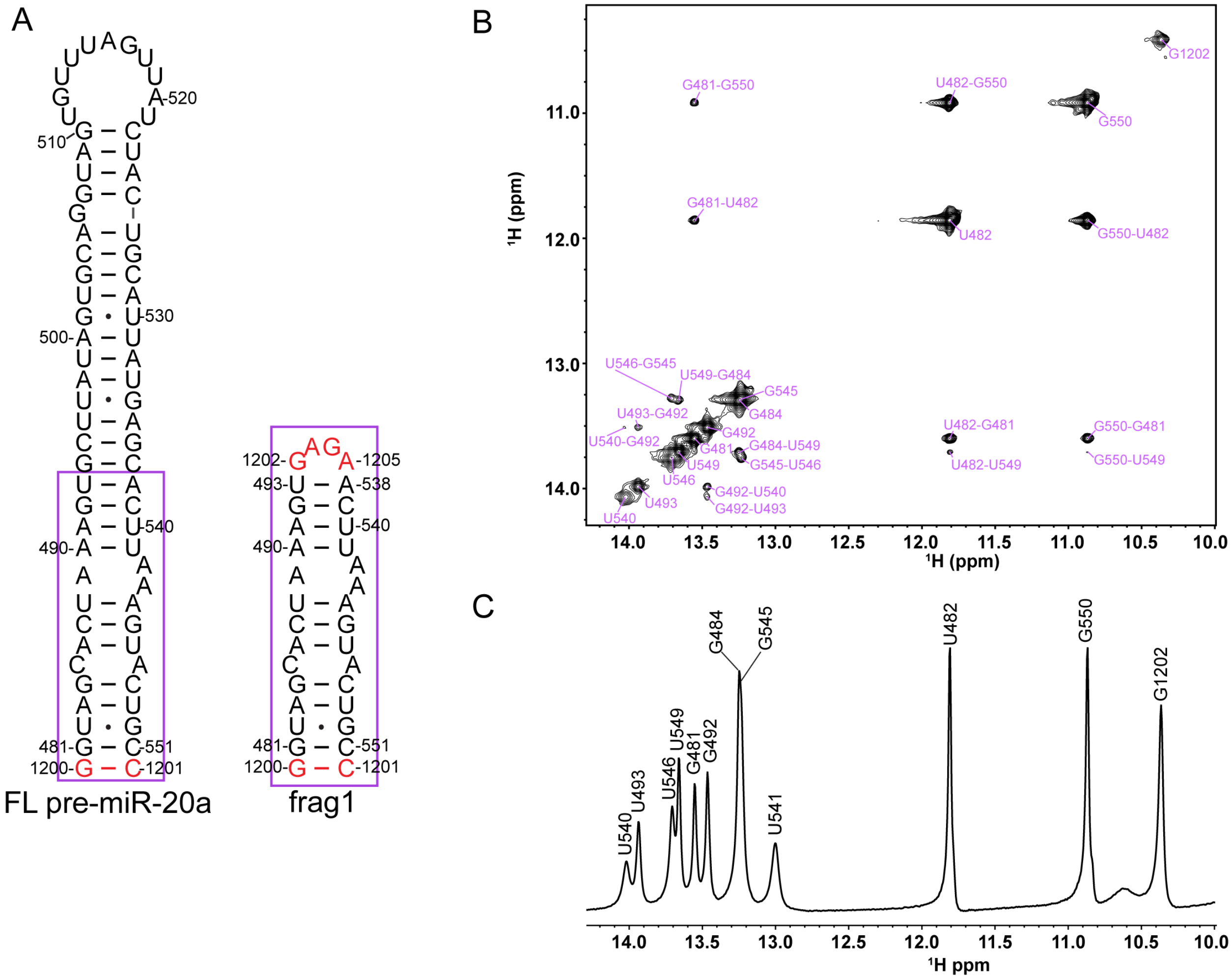

**Figure S10**. **Imino proton resonances of 20a-frag1.** (A) Secondary structure of FL pre-miR-20a and 20a-frag1. (B) Imino-imino region of 2D $^{1}$H-$^{1}$H NOESY spectrum of 20a-frag1. (C) 1D $^{1}$H NMR spectrum of 20a-frag1. The NMR spectra were recorded at 0.5 mM RNA concentration, 50 mM K-phosphate buffer, pH 6, 1 mM $MgCl_2$ and 90% $H_2O$/10% $D_2O$ at 600 MHz and at 15 °C.

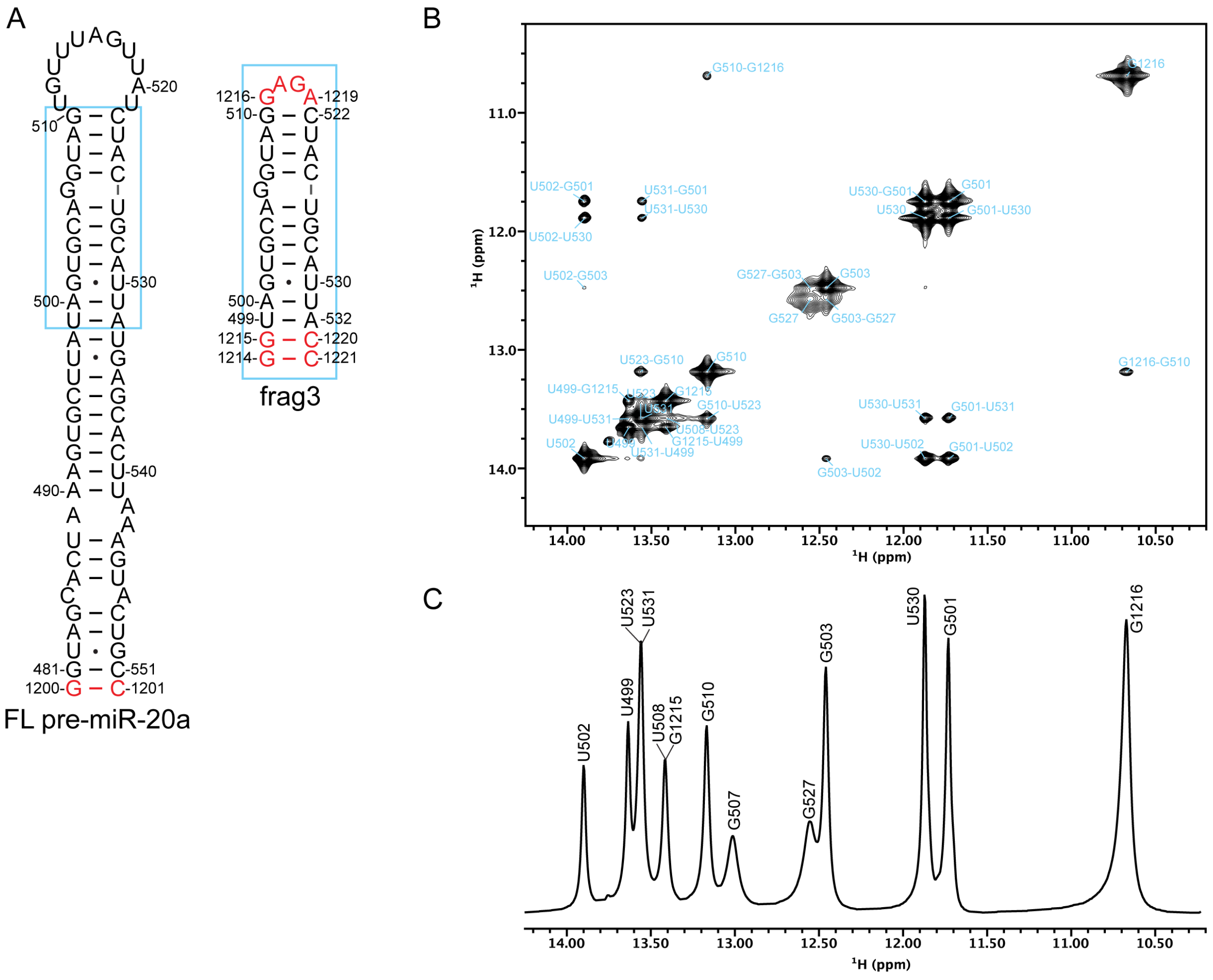

**Figure S11**. **Imino proton resonances of 20a-frag3.** (A) Secondary structure of FL pre-miR-20a and 20a-frag3. (B) Imino-imino region of 2D $^{1}H$-$^{1}H$ NOESY spectrum of 20a-frag3. (C) 1D $^{1}H$ NMR spectrum of 20a-frag3. The NMR spectra were recorded at 0.5 mM RNA concentration, 50 mM K-phosphate buffer, pH 6, 1 mM $MgCl_2$ and 90% $H_2O$/10% $D_2O$ at 600 MHz and at 0 °C.

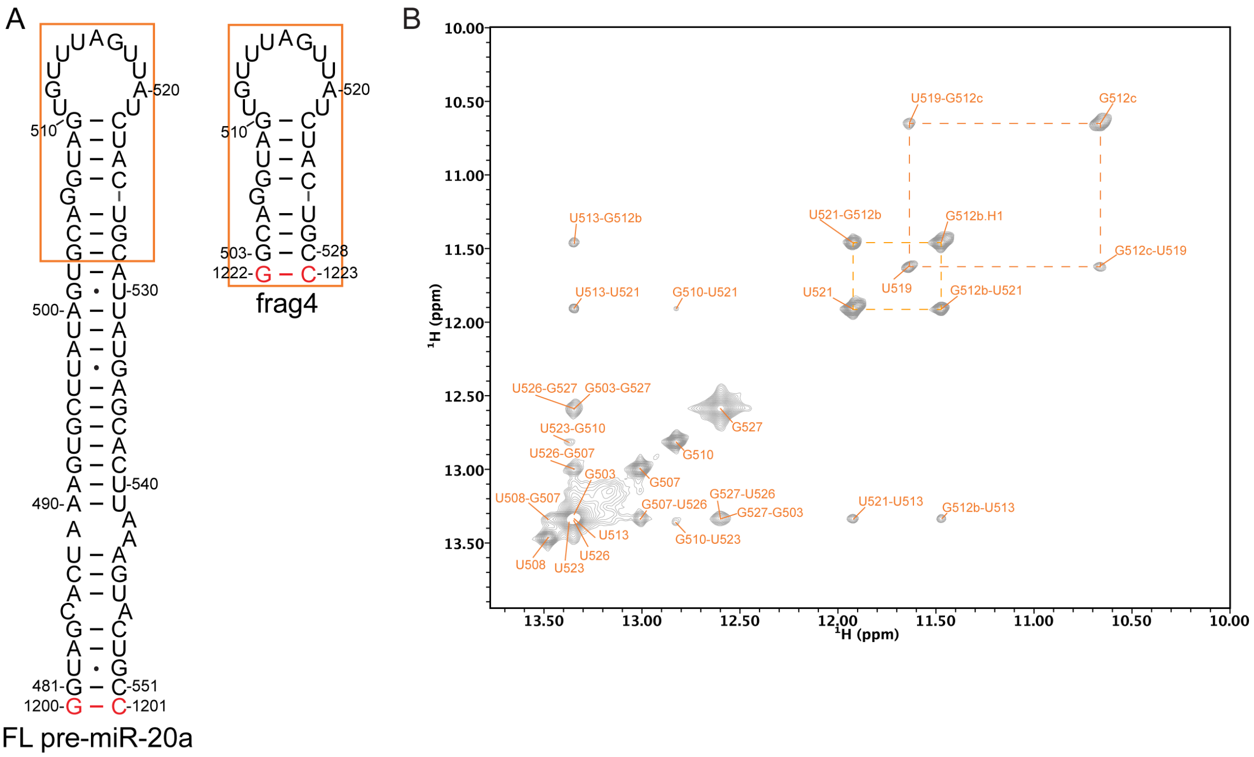

**Figure S12**. **Imino proton resonances of 20a-frag4.** (A) Secondary structure of FL pre-miR-20a and 20a-frag4. (B) Imino-imino region of 2D $^{1}H$-$^{1}H$ NOESY spectrum of 20a-frag4. The NMR spectra were recorded at 0.5 mM RNA concentration, 50 mM K-phosphate buffer, pH 6.5, 1 mM $MgCl_2$ and 90% $H_2O$/10% $D_2O$ at 600 MHz and at 10 °C.

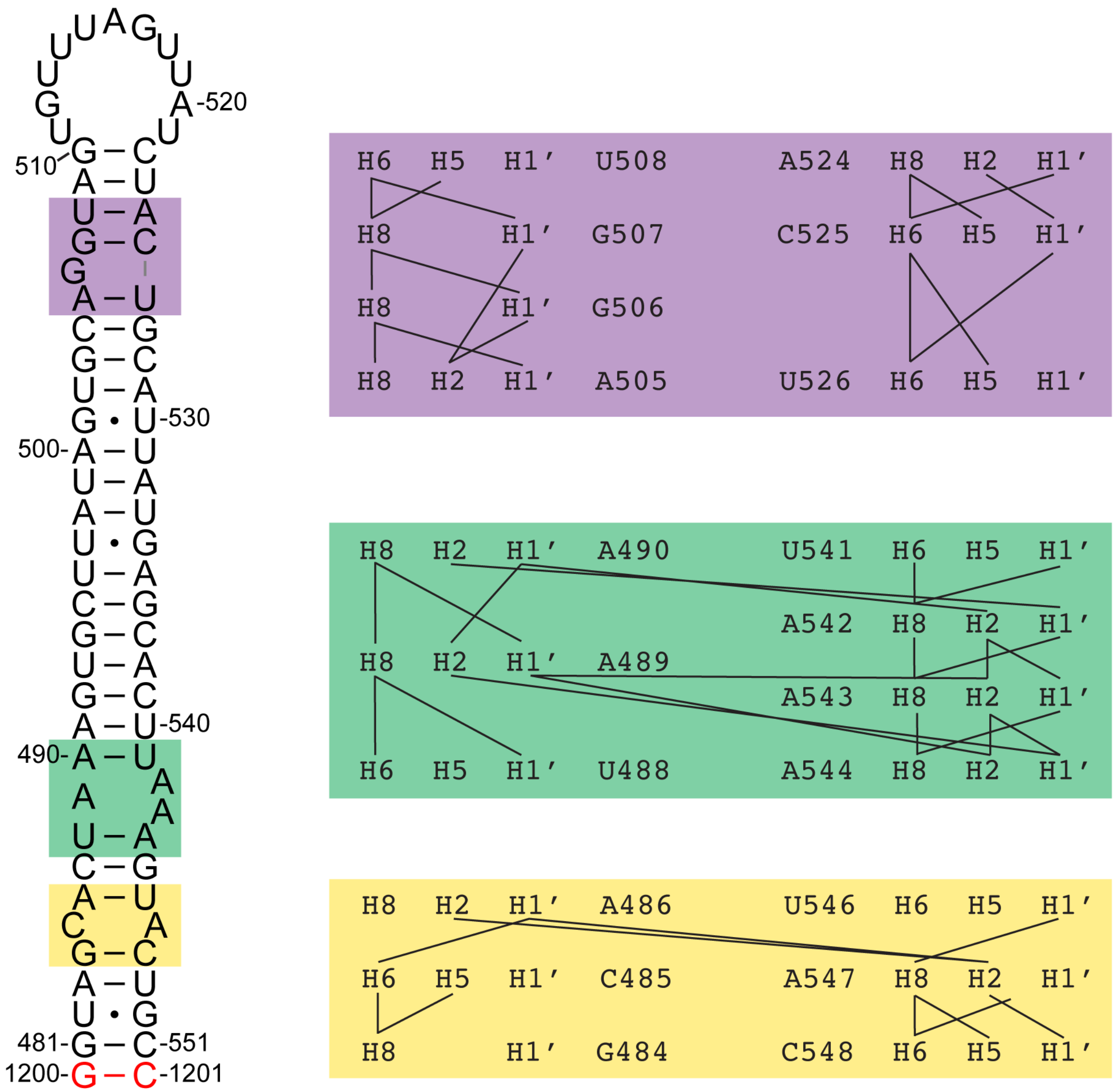

**Figure S13**. Summary of inter-residue NOEs for the G bulge (purple box), the A-rich 1x2 internal loop (green box), and the CA mismatch (yellow box).

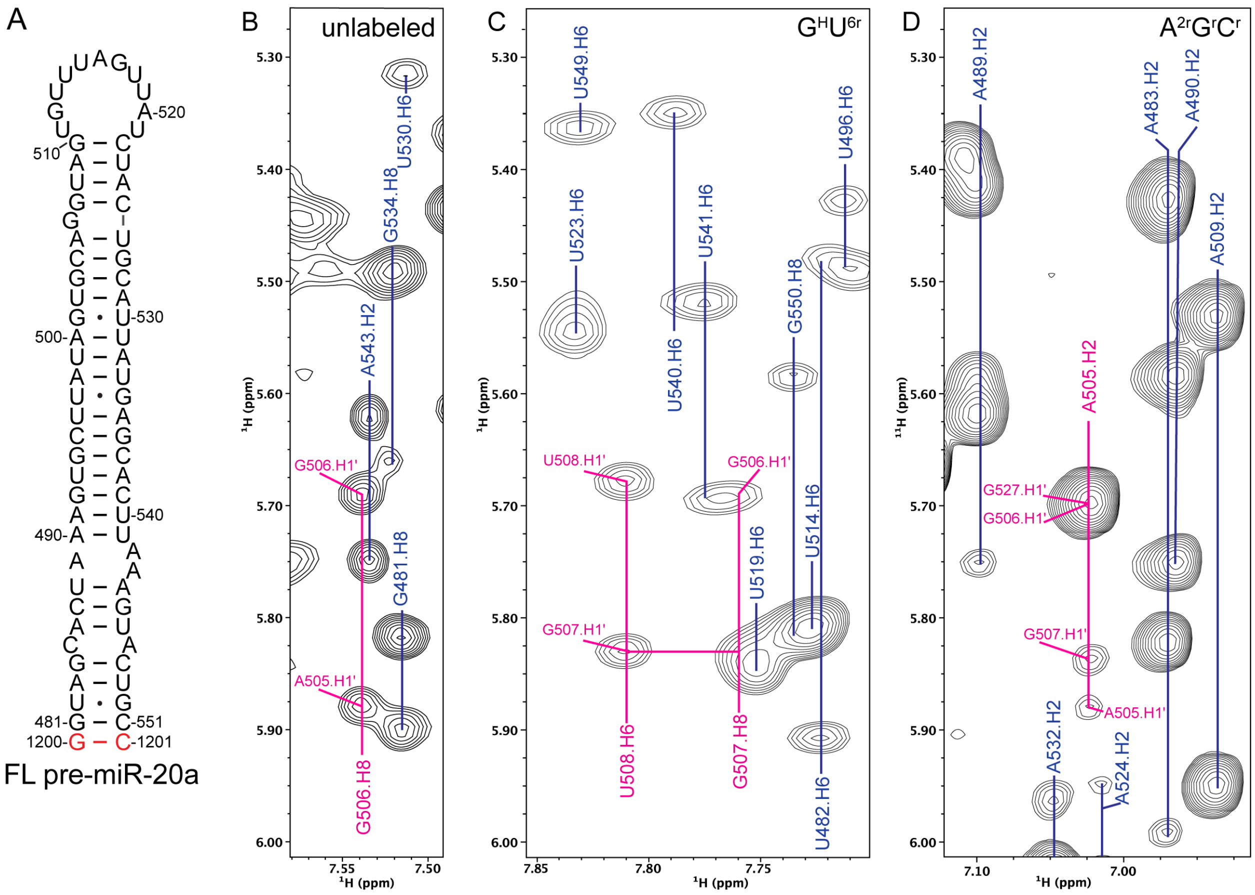

**Figure S14**. **NOEs of residues near the G bulge.** (A) Secondary structure of FL pre-miR-20a. (B) Region of the fully-protiated FL pre-miR-20a $^{1}$H-$^{1}$H NOESY spectrum, highlighting NOEs with G506.H8 in pink. Other assignments noted in blue. (C) Region of the $^{1}$H-$^{1}$H NOESY spectrum of $G^{H}U^{6r}$ labeled FL pre-miR-20a, highlighting NOEs connecting G507.H8 and U508.H6 in pink. Other assignments noted in blue. (D) Region of the $^{1}$H-$^{1}$H NOESY spectrum of $A^{2r}G^{r}C^{r}$ labeled FL pre-miR-20a, highlighting NOEs with A505.H2 in pink. Other assignments noted in blue.

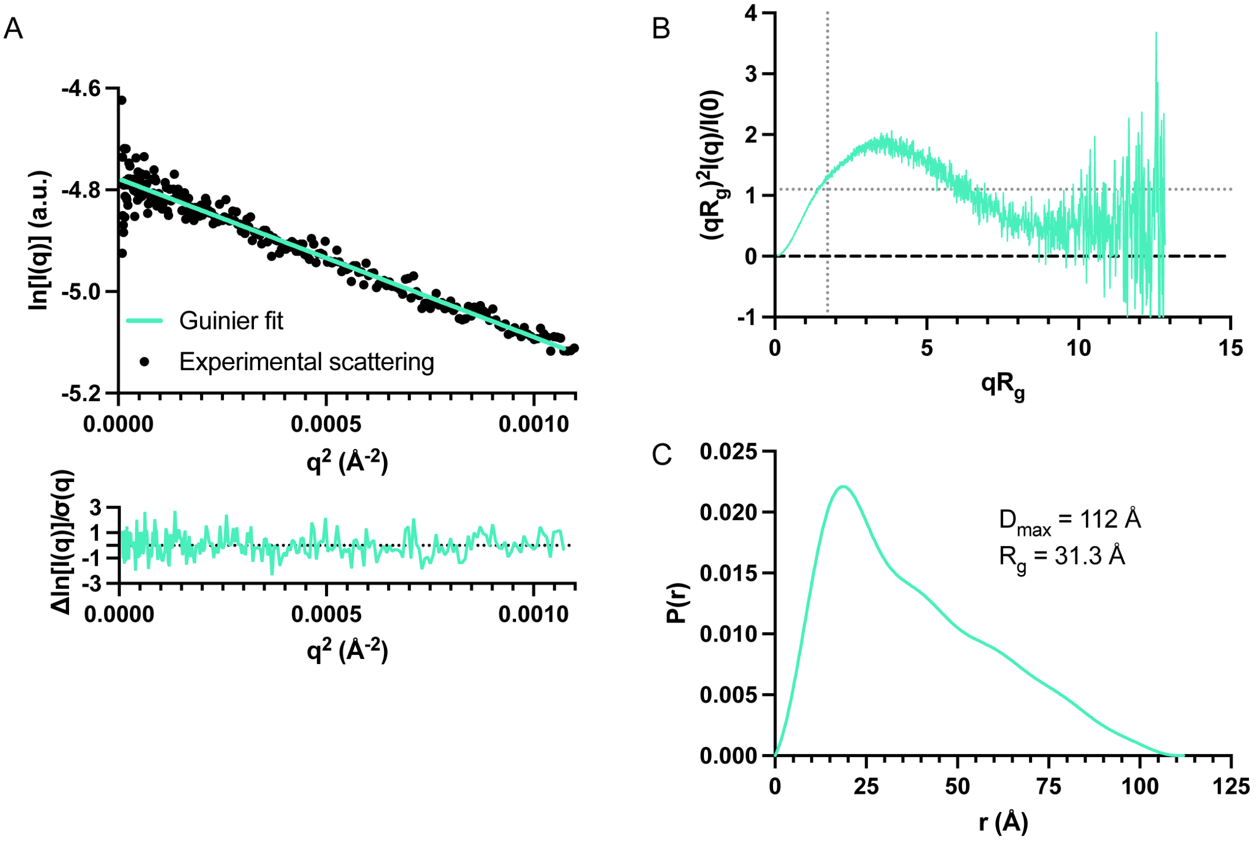

**Figure S15**. **Experimental SAXS data of FL pre-miR-20a.** (A) Guinier analysis of FL pre-miR-20a (top) with normalized residuals (bottom). (B) Dimensionless Kratky plot of FL pre-miR-20a. The dashed gray lines on the plot are guidelines for a peak position of a perfectly globular system. (C) Normalized pair distance distribution function of the FL pre-miR-20a.

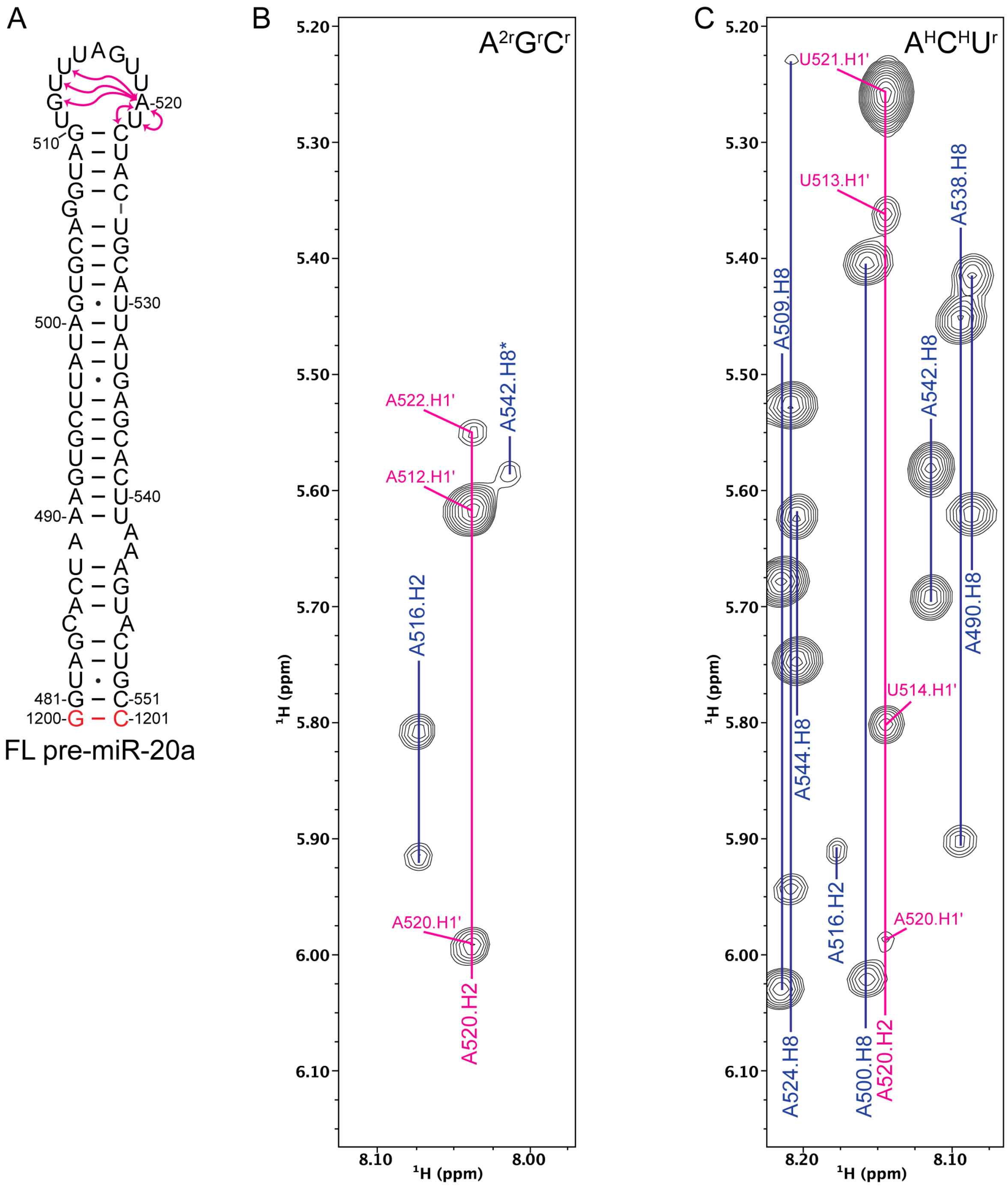

**Figure S16. Extensive cross-strand NOE network in the apical loop of FL pre-miR-20a.** (A) Secondary structure of FL pre-miR-20a highlighting extensive NOE connectivities (pink arrows). (B) Portion of 2D $^{1}$H-$^{1}$H NOESY spectrum of $A^{2r}G^{r}C^{r}$-labeled FL pre-miR-20a. NOEs with A520.H2 are labeled in pink. Other assignments are noted in blue. The * on A542.H8 indicates a breakthrough signal due to incomplete deuteration at the adenosine C8 position. (C) Portion of 2D $^{1}$H-$^{1}$H NOESY spectrum of $A^{H}C^{H}U^{r}$-labeled FL pre-miR-20a. NOEs with A520.H2 are labeled in pink. Other assignments are noted in blue.

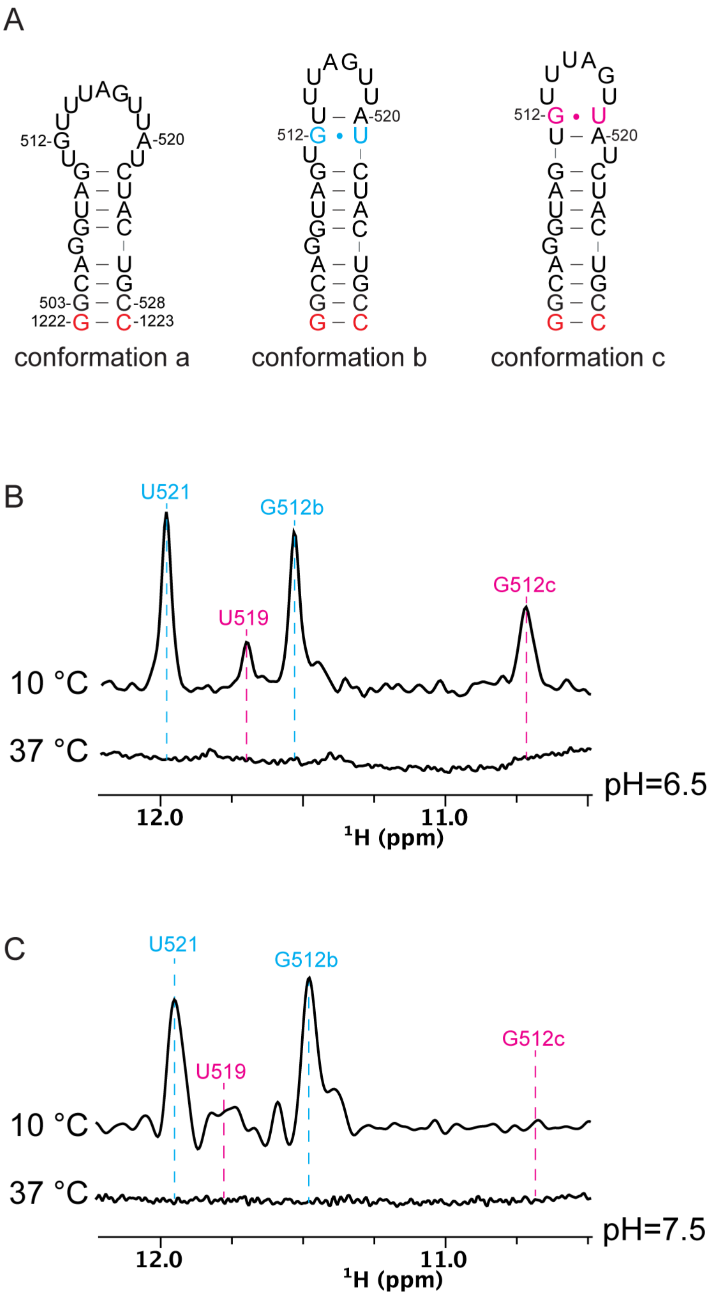

**Figure S17. Temperature-dependence of 20a-frag4 imino proton resonances.** (A) Secondary structures of alternative apical loop conformations. (B) Imino proton spectra of 20a-frag4 at pH 6.5. (C) Imino proton spectra of 20a-frag4 at pH 7.5. The NMR spectra were recorded at 0.5 mM RNA concentration, 50 mM K-phosphate buffer, 1 mM $MgCl_2$ and 90% $H_2O$/10% $D_2O$ at 600 MHz and at the temperature and pH indicated within the figure.

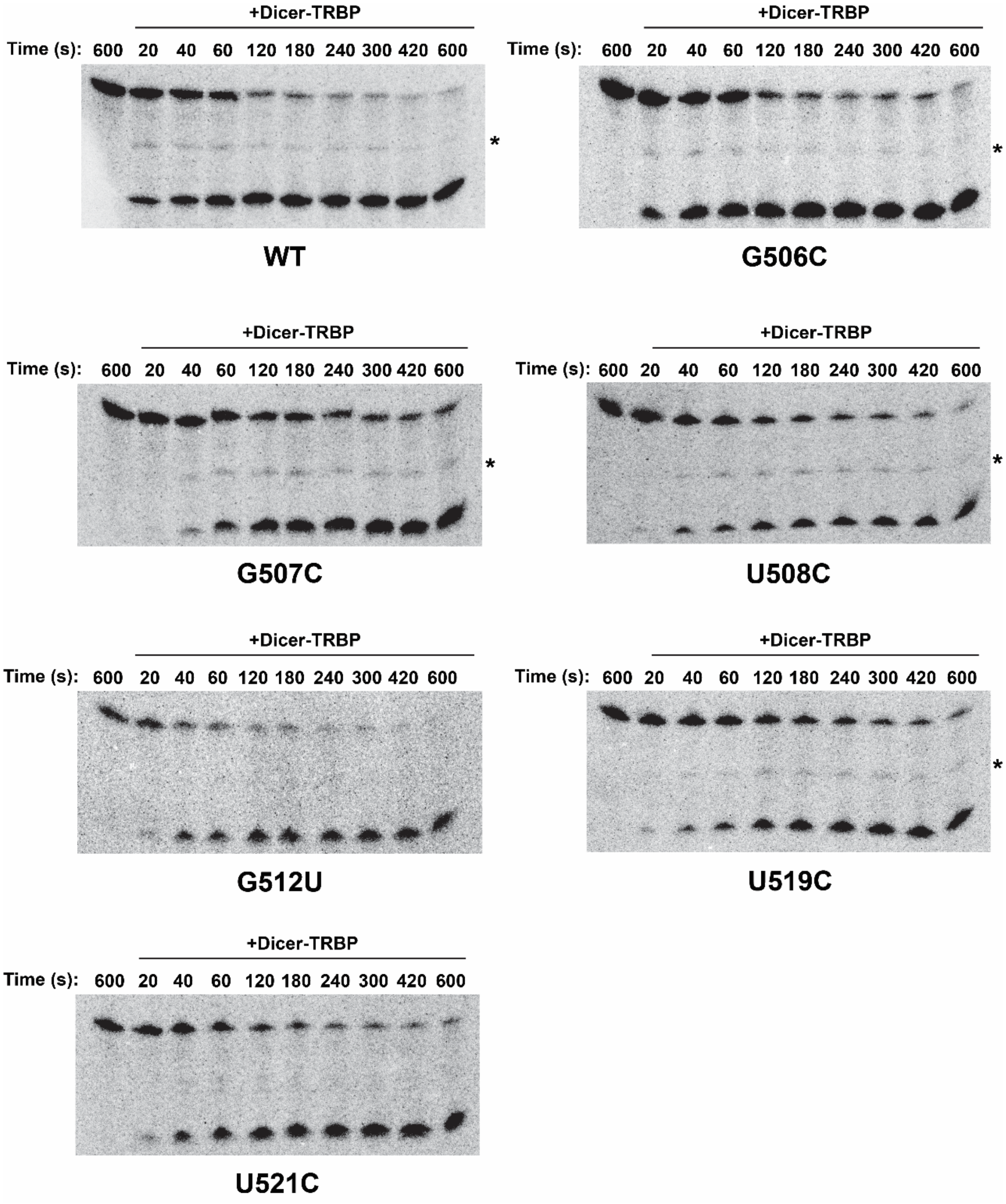

**Figure S18. Representative gel images from *in vitro* Dicer-TRBP processing assays.** *In vitro* Dicer-TRBP processing of $^{32}$P-labeled pre-miR-20a WT and mutant RNAs. The asterisk (*) indicates a partially processed product which was included in the total RNA calculation when present.

**Table S1.** Chemical shift completeness.[a]

| 20a-frag1 | A (% assigned) | C (% assigned) | G (% assigned) | U (% assigned) |
|---|---|---|---|---|
| H8/H6 | 100 | 100 | 100 | 100 |
| H2/H5 | 100 | 100 | / | 100 |
| H1′ | 100 | 100 | 100 | 100 |
| H2′ | 100 | 100 | 100 | 100 |
| H3′ | 91.7 | 100 | 100 | 100 |
| C6/C8 | 100 | 100 | 100 | 100 |
| C2 | 100 | / | / | / |
| 20a-frag2 | A (% assigned) | C (% assigned) | G (% assigned) | U (% assigned) |
| H8/H6 | 100 | 100 | 100 | 100 |
| H2/H5 | 100 | 100 | / | 100 |
| H1′ | 100 | 100 | 100 | 100 |
| H2′ | 100 | 100 | 100 | 100 |
| H3′ | 100 | 100 | 100 | 100 |
| C6/C8 | 100 | 100 | 100 | 100 |
| C2 | 100 | / | / | / |
| 20a-frag3 | A (% assigned) | C (% assigned) | G (% assigned) | U (% assigned) |
| H8/H6 | 100 | 100 | 100 | 100 |
| H2/H5 | 100 | 100 | / | 100 |
| H1′ | 100 | 100 | 100 | 100 |
| H2′ | 100 | 100 | 100 | 100 |
| H3′ | 100 | 100 | 100 | 100 |
| C6/C8 | 100 | 100 | 100 | 100 |
| C2 | 100 | / | / | / |
| 20a-frag4 | A (% assigned) | C (% assigned) | G (% assigned) | U (% assigned) |
| H8/H6 | 100 | 100 | 100 | 100 |
| H2/H5 | 100 | 100 | / | 100 |
| H1′ | 100 | 100 | 100 | 100 |
| H2′ | 100 | 100 | 100 | 100 |
| H3′ | 100 | 100 | 100 | 100 |
| C6/C8 | 100 | 100 | 100 | 100 |
| C2 | 100 | / | / | / |
| FL pre-miR-20a | A (% assigned) | C (% assigned) | G (% assigned) | U (% assigned) |
| H8/H6 | 100 | 100 | 100 | 100 |
| H2/H5 | 100 | 100 | / | 100 |
| H1′ | 100 | 100 | 100 | 100 |
| H2′ | 100 | 100 | 100 | 100 |
| H3′ | 100 | 100 | 100 | 100 |
| C6/C8 | 100 | 100 | 100 | 100 |
| C2 | 100 | / | / | / |

[a] "/" indicates a given atom is not present in the nucleoside.

**Table S2.** NMR restraints and structural statics for the FL pre-miR-20a structure.[a]

| | |
|---|---|
| **CYANA[b]** | |
| NOE-derived restraints | 390 |
| Intraresidue | 147 |
| Sequential | 205 |
| Long-range (\|i-j\| > 1) | 38 |
| H-bond restraints | 268 |
| NOE restraints/residue | 5.3 |
| Target function (Å$^2$) | 0.118 ± 0.0250 |
| Upper distance viol. (Å$^2$) | 0.00837 ± 0.000867 |
| Lower distance viol. (Å$^2$) | 0.00502 ± 0.000834 |
| RMSD[c] (Å) | 1.15 ± 0.344 |
| **Amber[d]** | |
| Amber Energy | -16,532 |
| Distance | 4.688 |
| Torsion | 35.504 |
| RMSD[c] (Å) | 1.00 ± 0.308 |
| **MolProbity analysis[e]** | |
| Clashscore | 0.86 |
| Probably wrong sugar pucker (%) | 0 |
| Bad backbone conformation (%) | 6.5 |
| Bad bonds (%) | 0 |
| Bad angles (%) | 0 |

[a]Statistics are reported for the entire structure, unless otherwise indicated.
[b]Statistics for the 20 structures with lowest target function.
[c]Root mean square deviation. Statistics are reported for residues 481-510, 522-551
[d]Statistics for the 20 lowest energy structures.
[e]The 20 Amber-refined structures were evaluated using the MolProbity webserver.[1,2]

**Table S3.** SEC-SAXS data acquisition, sample details, data analysis, model fitting, and software used.

| | |
|---|---|
| **(a) Sample details** | |
| Sample | FL pre-miR-20a |
| Organism | *Homo sapiens* |
| Sequence (5′ to 3′) | GGUAGCACUAAAGUGCUUAUAGUGCAGGUAGUGUUUAGUUAUCUACUGCAUUAUGA GCACUUAAAGUACUGCC |
| Extinction coefficient, $\varepsilon_{260}$ ($M^{-1}$ $cm^{-1}$) | 744800 |
| *M* from chemical composition (kDa) | 23.48 |
| SEC-SAXS column | Superdex 75 10/300 Increase |
| Loading volume (μL) | 260 |
| Concentration (mg/mL) | 1.3 |
| Flow rate (mL/min) | 0.6 |
| Solvent | 50 mM potassium phosphate buffer pH = 7.5, 1 mM $MgCl_2$, 50 mM NaCl, 5 mM BME |
| **(b) SAS data collection parameters** | |
| Instrument | BioCAT (beamline 18ID, APS) with a Dectris Eiger2 XE 9M detector |
| Wavelength (Å) | 1.033 |
| Beam size (μm) | 30 × 150 (focused on detector) |
| Camera length (m) | 3.663 |
| $q$ measurement range ($Å^{-1}$) | 0.0029 – 0.42 |
| Absolute scaling method | Glassy Carbon, NIST SRM 3600 |
| Normalization | To transmitted intensity by beam-stop counter |
| Monitoring for radiation damage | Automated frame-by-frame comparison of relevant regions using CORMAP[3] implemented in BioXTAS RAW[4] |
| Exposure time | 0.5 s exposure time with a 1 s total exposure period (0.5 s on, 0.5 s off) of entire SEC elution |
| Sample configuration | SEC-MALS-SAXS. Size separation used a Superdex 75 10/300 Increase column and a 1260 Infinity II HPLC (Agilent Technologies). UV data was measured in the Agilent, and MALS-DLS-RI data by DAWN HELEOS-II (17 MALS + 1 DLS channels) and Optilab T-rEX (RI) instruments (Wyatt Technology). SAXS data was measured in a sheath-flow cell,[5] effective path length 0.542 mm. |
| Sample temperature (°C) | 22 |
| **(c) Software employed for SAXS data reduction, analysis, and interpretation** | |
| Data reduction | $I(q)$ vs $q$ and solvent subtraction using BioXTAS RAW 2.2.1[4] |
| Extinction coefficient estimate | Quest Calculate™ RNA Concentration Calculator *via* web server (https://www.aatbio.com/tools/calculate-RNA concentration) |

| | |
|---|---|
| Basic analyses, Guinier, $P(r)$, $V_p$ | BioXTAS RAW 2.2.1[6] and GNOM from ATSAS 3.0.307/05/2025 13:52:00 |
| Electron density modelling | DENSS[3] |
| Atomic structure modelling | FoXS[7,8] *via* web server (https://modbase.compbio.ucsf.edu/foxs/) |
| Three-dimensional graphic model representations | PyMOL (version 3.1.4.1) |
| **(d) Structural parameters** | |
| Guinier analysis | |
| $I(0)$ ($cm^{-1}$) | $0.01 \pm 1.86\times10^{-5}$ |
| $R_g$ (Å) | $30.57 \pm 0.19$ |
| $q_{min}$ ($Å^{-1}$) | 0.0029 |
| $qR_g$ max | 1.001 |
| Coefficient of correlation, $R^2$ | 0.936 |
| $M$ from $V_c$ | 25.2 |
| $P(r)$ analysis | |
| $I(0)$ ($cm^{-1}$) | $0.01 \pm 1.75\times10^{-5}$ |
| $R_g$ (Å) | $31.29 \pm 0.13$ |
| $D_{max}$ (Å) | 112 |
| $q$-range ($Å^{-1}$) | 0.0029 – 0.4201 |
| $\chi^2$ | 0.872 |
| **(e) Shape model-fitting results** | |
| DENSS (default parameters, 20 calculations) | |
| $q$-range for fitting | 0.0029 – 0.4201 |
| Symmetry, anisotropy assumptions | P1, none |
| Ambiguity score (AMBIMETER)[9] | 2.155 |
| $\chi^2$ range | 0.00283 - 0.06927 |
| Model resolution (Å) | $25.4 \pm 3.9$ |
| **(f) Atomistic modeling** | |
| NMR structures | PDB entry 9OBM |
| FoXS | |
| $\chi^2$ | 0.88 |
| Predicted $R_g$ (Å) | 30.97 |
| $c_1$, $c_2$ | 1.01, 2.97 |
| **(g) SASBDB IDs for data and models** | |
| FL pre-miR-20a | SASXXXX |

**Table S4.** Synthetic DNA templates and associated RNA constructs.

| | 5′-sequence-3′[a,b,c] | |
|---|---|---|
| Construct | DNA | RNA |
| 20a-frag1 | mGmGCAGTACTTTAAGTTCTCACTTTAGTGC TACC*TATAGTGAGTCGTATTA* | GGUAGCACUAAAGUGAGAACUUAA AGUACUGCC |
| 20a-frag2 | mGmGAAGTGCTCATAATTCTCACTATAAGC ACTTCC*TATAGTGAGTCGTATTA* | GGAAGUGCUUAUAGUGAGAAUUA UGAGCACUUCC |
| 20a-frag3 | mGmGTAATGCAGTAGTCTCCTACCTGCACT ACC*TATAGTGAGTCGTATTA* | GGUAGUGCAGGUAGGAGACUACUG CAUUACC |
| 20a-frag4 | mGmGCAGTAGATAACTAAACACTACCTGCC *TATAGGAGTCGTATTA* | GGCAGGUAGUGUUUAGUUAUCUAC UGCC |

[a] m denotes 2′-*O*-Me modification of the primer.

[b] Italicized nucleotides correspond to the sequence complementary to the T7 promoter.

[c] Red nucleotides indicate non-native sequences.

**Table S5.** DNA primers for generation of the FL pre-miR-20a (NMR/SAXS) template.

| | 5′-sequence-3′[a] |
|---|---|
| Pre-miR-20a_F | TTCTAATACGACTCACTATAGGTAGCACTAAAGTGCTTATAGTGCAGGTAGTGT |
| Pre-miR-20a_R | mGmGCAGTACTTTAAGTGCTCATAATGCAGTAGATAACTAAACACTACCTGCA CTATAAGCA |

[a] m denotes 2′-*O*-Me modification of the primer.

**Table S6.** DNA primers for HH-pre-miR-20a-HDV template.

| | 5′-sequence-3′ |
|---|---|
| HH-20a-HDV_F | CCGGAATTCTAATACGACTCACTATAGGGCTCG |
| HH-20a-HDV_R | CCGTCGCGGATCCTAATGTGAGAATTGGCTACGTTGA |

**Table S7.** Mutation DNA primers for processing constructs.

| | 5′-sequence-3′ | Application |
|---|---|---|
| 20a-G506C_F | TAGTGTTTAGTTATCTACTGCATTATG | Forward primer for G506C mutation |
| 20a-G506C_R | CGTGCACTATAAGCACTTTAGACG | Reverse primer for G506C mutation |
| 20a-G507C_F | AGTGTTTAGTTATCTACTGCATTATG | Forward primer for G507C mutation |
| 20a-G507C_R | AGCTGCACTATAAGCACTTTAGAC | Reverse primer for G507C mutation |
| 20a-U508C_F | ATAGTGCAGGCAGTGTTTAGTTATC | Forward primer for U508C mutation |
| 20a-U508C_R | AAGCACTTTAGACGGTACCGG | Reverse primer for U508C mutation |
| 20a-G512U_F | TGCAGGTAGTTTTTAGTTATCTAC | Forward primer for G512U mutation |
| 20a-G512U_R | CTATAAGCACTTTAGACG | Reverse primer for G512U mutation |
| 20a-U519C_F | AGTGTTTAGTCATCTACTGCATTATG | Forward primer for U519C mutation |
| 20a-U519C_R | ACCTGCACTATAAGCACTTTAG | Reverse primer for U519C mutation |
| 20a-U521C_F | TGTTTAGTTACCTACTGCATTATG | Forward primer for U521C mutation |
| 20a-U521C_R | CTACCTGCACTATAAGCAC | Reverse primer for U521C mutation |

**Table S8.** Amplification primers for template.

| Amplification primers | 5′-sequence-3′[a] | Application |
|---|---|---|
| UNIV-pUC19_E105 | TCTTCGCTATTACGCCAGCTGGCGAAA | Forward primers for amplification of DNA templates for all pre-miR-20a processing constructs |
| HDV-AMP-R | mUmAATGTGAGAATTGGCTACGTTGA AACAACGCATTACCG | Reverse primers for amplification of DNA templates for all pre-miR-20a processing constructs |

[a] m denotes 2′-*O*-Me modification of the primer.

**Table S9.** RNA sequence used for structural and processing studies.

| Construct | 5′-sequence-3′ | Application |
|---|---|---|
| FL pre-miR-20a | GGUAGCACUAAAGUGCUUAUAGUGCAGGUAGUGUUU AGUUAUCUACUGCAUUAUGAGCACUUAAAGUACUGCC | NMR/SAXS studies |
| pre-miR-20a | UAAAGUGCUUAUAGUGCAGGUAGUGUUUAGUUAUCU ACUGCAUUAUGAGCACUUAAAG | Processing |
| pre-miR-20a_G506C | UAAAGUGCUUAUAGUGCACGUAGUGUUUAGUUAUCU ACUGCAUUAUGAGCACUUAAAG | Processing |
| pre-miR-20a_G507C | UAAAGUGCUUAUAGUGCAGCUAGUGUUUAGUUAUCU ACUGCAUUAUGAGCACUUAAAG | Processing |
| pre-miR-20a_U508C | UAAAGUGCUUAUAGUGCAGGCAGUGUUUAGUUAUCU ACUGCAUUAUGAGCACUUAAAG | Processing |
| pre-miR-20a_G512U | UAAAGUGCUUAUAGUGCAGGUAGUUUUUAGUUAUCU ACUGCAUUAUGAGCACUUAAAG | Processing |
| pre-miR-20a_U519C | UAAAGUGCUUAUAGUGCAGGUAGUGUUUAGUCAUCU ACUGCAUUAUGAGCACUUAAAG | Processing |
| pre-miR-20a_U521C | UAAAGUGCUUAUAGUGCAGGUAGUGUUUAGUUACCU ACUGCAUUAUGAGCACUUAAAG | Processing |

**Table S10.** Dicer-TRBP processing of pre-miR-20a RNAs.[a]

| RNA construct | % full cleavage (10 min.) | $k_{obs}$ ($s^{-1}$) | $k_{obs}$ fold-change rel. to WT[b] | Adjusted p-value rel. to WT[c] | Significance level[c] |
|---|---|---|---|---|---|
| WT | 97.0 ± 0.9 | 0.01581 ± 0.00101 | ---- | ---- | ---- |
| G506C | 97.1 ± 1.3 | 0.01450 ± 0.00029 | 0.92 | 0.2825 | ns |
| G507C | 83.6 ± 0.3 | 0.00893 ± 0.00107 | 0.56 | <0.0001 | **** |
| U508C | 88.2 ± 2.2 | 0.00484 ± 0.00047 | 0.31 | <0.0001 | **** |
| G512U | 98.0 ± 2.2 | 0.01838 ± 0.00075 | 1.16 | 0.0116 | * |
| U519C | 86.5 ± 0.7 | 0.00572 ± 0.00057 | 0.36 | <0.0001 | **** |
| U521C | 92.5 ± 1.6 | 0.01196 ± 0.00117 | 0.76 | 0.0004 | *** |

[a]Values represent average and standard deviation from n = 2-3 independent assays.

[b]$k_{obs}(mut)/k_{obs}(WT)$

[c]Adjusted p-values are from one-way ANOVA with Dunnett's multiple comparison test (relative to mean of WT, $\alpha = 0.05$). * $p < 0.05$, ** $p < 0.01$, *** $p < 0.001$, **** $p < 0.0001$.
